# TopFor: Automated Preparation of AMBER-Compatible Parameters for Noncanonical Amino Acids in Peptides

**DOI:** 10.64898/2026.10.05.756260

**Authors:** Emre F. Bülbül, Heshine Gnanasekar, Carène A. Benasolo, Sari Rasheed, Rolf Müller, Olga V. Kalinina, Kenan A. J. Bozhüyük

**Affiliations:** Helmholtz Institute for Pharmaceutical Research Saarland (HIPS), Helmholtz Centre for Infection Research (HZI), PharmaScienceHub (PSH), Campus E8.1, 66123, Saarbrücken, Germany; Center for Bioinformatics, Saarland University, Campus E2.1, 66123, Saarbrücken, Germany; German Centre for Infection Research (DZIF), Partner Site Hannover-Braunschweig, Germany; Department of Pharmacy, Saarland University, Campus E8.1, 66123, Saarbrücken, Germany; Medical Faculty, Saarland University, 66421, Homburg, Germany; Myria Biosciences AG, Hochbergerstrasse 60C, 4057, Basel, Switzerland

**Author notes:** These authors contributed equally (Emre F. Bülbül, Heshine Gnanasekar).

**Keywords:** noncanonical amino acids, peptide parameterization, molecular dynamics, AMBER, AmberTools, peptide design, high-throughput virtual screening

## Abstract

Peptides containing noncanonical amino acids (ncAAs) are becoming increasingly important in drug discovery, but their molecular dynamics (MD) simulation remains limited by labor-intensive residue parameterization and topology preparation. Here, we present TopFor (Topology and Residue Force-Field Parameter Preparation), an open-source, peptide-oriented workflow that automates the generation of ncAA residue templates and force-field parameter files for MD simulations with AMBER using established charge-derivation methods. TopFor supports single-residue, whole-peptide, and batch parameterization workflows.

The workflow was validated using the MDM2–p53 complex by comparing native AMBER residue definitions with residues processed by TopFor using three charge-derivation methods, each evaluated in three independent 100 ns MD simulations. TopFor-generated residue definitions showed structural and conformational behavior comparable to the native AMBER representation. Applicability to more complex peptide architectures was demonstrated using the head-to-tail cyclic peptide cyclosporin A and the cyclic depsipeptide cyclohexylgriselimycin. At library scale, 1,138 of 1,214 cyclohexylgriselimycin derivatives (93.7%) successfully completed TopFor parameterization and the subsequent initial AMBER-based computational workflow across two derivatization campaigns. TopFor was further integrated into a structure-based peptide discovery workflow based on the DnaN–griselimycin complex. Six GM-derived peptides were computationally prioritized and experimentally evaluated, of which four produced SPR responses against *Mycobacterium smegmatis (M. smegmatis)* DnaN and three showed concentration-dependent responses.

TopFor reduces the manual effort required to prepare ncAA-containing peptides for MD simulations and enables their integration into both library-scale computational screening and structure-based peptide discovery workflows.

## Introduction

Peptides have emerged as an important therapeutic modality that occupies a unique position between small molecules and biologics^1,2^. Owing to their larger interaction surfaces and high molecular recognition, peptides can bind to protein–protein interfaces and to other challenging biological targets that are often inaccessible to conventional small molecules while retaining greater synthetic accessibility than antibodies^3^. Since the introduction of the peptide hormone insulin nearly a century ago, more than 80 peptide therapeutics have been approved for the treatment of diseases including diabetes, cancer, infectious diseases, osteoporosis, multiple sclerosis, and chronic pain^4^. Despite these successes, peptide therapeutics remain underrepresented relative to their therapeutic potential because many peptide candidates exhibit rapid proteolytic degradation, limited membrane permeability, poor oral bioavailability, and short plasma half-lives^5^. These limitations frequently restrict peptide therapeutics to intravenous or subcutaneous administration^6^.

Peptide natural products, particularly nonribosomal peptides (NRPs) and ribosomally synthesized and post-translationally modified peptides (RiPPs), provide examples of diverse chemical modifications, including macrocyclization, *N*-methylation, incorporation of D- and other noncanonical amino acids (ncAAs), and heterocyclization^7–10^. These structural features expand the chemical and conformational space accessible to peptides and can modulate their stability, membrane permeability, target affinity, and pharmacological properties^11^. Cyclosporin A, for example, is a head-to-tail cyclic peptide containing nine noncanonical amino acids and combines potent immunosuppressive activity with oral bioavailability^12–14^. Another example is cyclohexylgriselimycin (CGM)^15,16^, a cyclic depsipeptide containing seven ncAA building blocks. Inspired by such natural products, modern peptide engineering exploits ncAAs to expand accessible chemical space and optimize the properties of peptide candidates^6,10,17–20^.

Although peptide generation has become accessible through experimental screening, synthetic chemistry, synthetic biology, and AI-based design^21–26^, computational evaluation of the resulting candidates remains challenging. Molecular dynamics (MD) simulations provide an atomistic framework for investigating peptide conformational behavior and interactions with molecular targets. However, MD simulations of peptides containing ncAAs require force-field parameters for residues that are not generally covered by standard protein force fields.

Several computational tools facilitate molecular parameterization, but each addresses only part of the workflow required for peptides containing ncAAs. General small-molecule tools such as ACPYPE automate topology generation through the Antechamber/GAFF workflow for multiple simulation packages but are designed for isolated ligands rather than amino acid residues incorporated into peptide chains^27^. Within the CHARMM ecosystem, the CHARMM36^28^/CHARMM General force field^29^ (CGenFF) has been systematically extended by developing parameters for a predefined set of 333 ncAAs, including residues with modified side chains and nonstandard backbone groups^29,30^. This work provides a comprehensive library of validated parameters for the included residues, generated through extensive QM-based optimization of partial charges and bonded parameters. However, parameterization of a new ncAA outside the available set still requires force-field development, including QM calculations and parameter optimization^29,30^. More recently, dedicated ncAA workflows have emerged. NCAP automatically generates CHARMM-compatible parameters for amino acid derivatives but is restricted to the CHARMM force-field ecosystem and relies on quantum mechanical calculations^31^. Similarly, drFrankenstein provides an automated AMBER workflow incorporating terminal capping, restrained electrostatic potential (RESP) charge derivation, and parameter fitting, but requires multiple quantum mechanical calculations for each residue and is primarily intended for individual rather than library-scale parameterization^32^. Wijesiri and Brown described a protocol for generating ncAA residue definitions for Rosetta modeling and AMBER MD, including conformational sampling, QM calculations, multi-conformer RESP fitting, and generation of AMBER residue templates and force-field parameters^33^. More recently, Pep2MARS^34^ was introduced as an automated AMBER-compatible workflow for parameterizing cyclic peptides containing noncanonical amino acids and complex covalent linkages through connectivity-based identification and parameterization of covalently connected residue blocks. The reported applications demonstrate automated parameterization across diverse cyclization patterns, whereas library-scale parameterization of large numbers of peptide derivatives was not evaluated.

Nevertheless, preparation of large libraries of ncAA-containing peptides for MD simulations remains challenging. Existing approaches differ in molecular scope, force-field ecosystem, computational requirements, and degree of peptide-specific automation, which can limit their application to large residue libraries. In addition, combining automatic identification of ncAAs from complete peptide structures, recognition of their position within the peptide chain, batch parameterization, and generation of residue templates and force-field parameter files for system assembly within a single workflow remains challenging. Consequently, preparation of large ncAA-containing peptide libraries for MD simulations can require substantial manual intervention, limiting throughput and reproducibility.

Among available MD simulation frameworks, AMBER provides an established ecosystem for biomolecular simulations and residue parameterization through AmberTools^35,36^. We therefore developed TopFor (Topology and Residue Force-Field Parameter Preparation) within this framework to automate the generation of residue templates and force-field parameter files for peptides containing ncAAs. Rather than introducing new force-field parameters or charge models, TopFor automates residue-parameterization procedures within a peptide-oriented workflow. The workflow supports single-residue, batch, and whole-peptide processing and multiple charge-derivation methods, including AM1-BCC^37,38^, ABCG2^39,40^, RESP^41^, and Gasteiger^42^. We evaluate the workflow using linear, head-to-tail cyclic, and cyclic depsipeptide systems, assess its scalability in two high-throughput peptide derivatization campaigns, and demonstrate the use of TopFor-generated parameters within a structure-based peptide discovery workflow.

## Methods

### TopFor Workflow Overview

TopFor is an automated command-line workflow (Figure 1) for preparing AMBER^36^-compatible residue topologies and force-field parameter files for peptides containing ncAAs. The generated files can be loaded into the LEaP module of AmberTools^35^ together with the selected force field for peptide system assembly. TopFor supports three input modes: (i) a single ncAA residue, (ii) batch mode for parameterization of multiple ncAA residue structures, and (iii) whole-peptide mode for processing a complete peptide structure. In whole-peptide mode, TopFor identifies ncAAs within the input structure, determines their position within the peptide chain (*N*-terminal, *C*-terminal, or internal), extracts the corresponding residues, and processes each ncAA according to its residue context.

**Figure 1.**
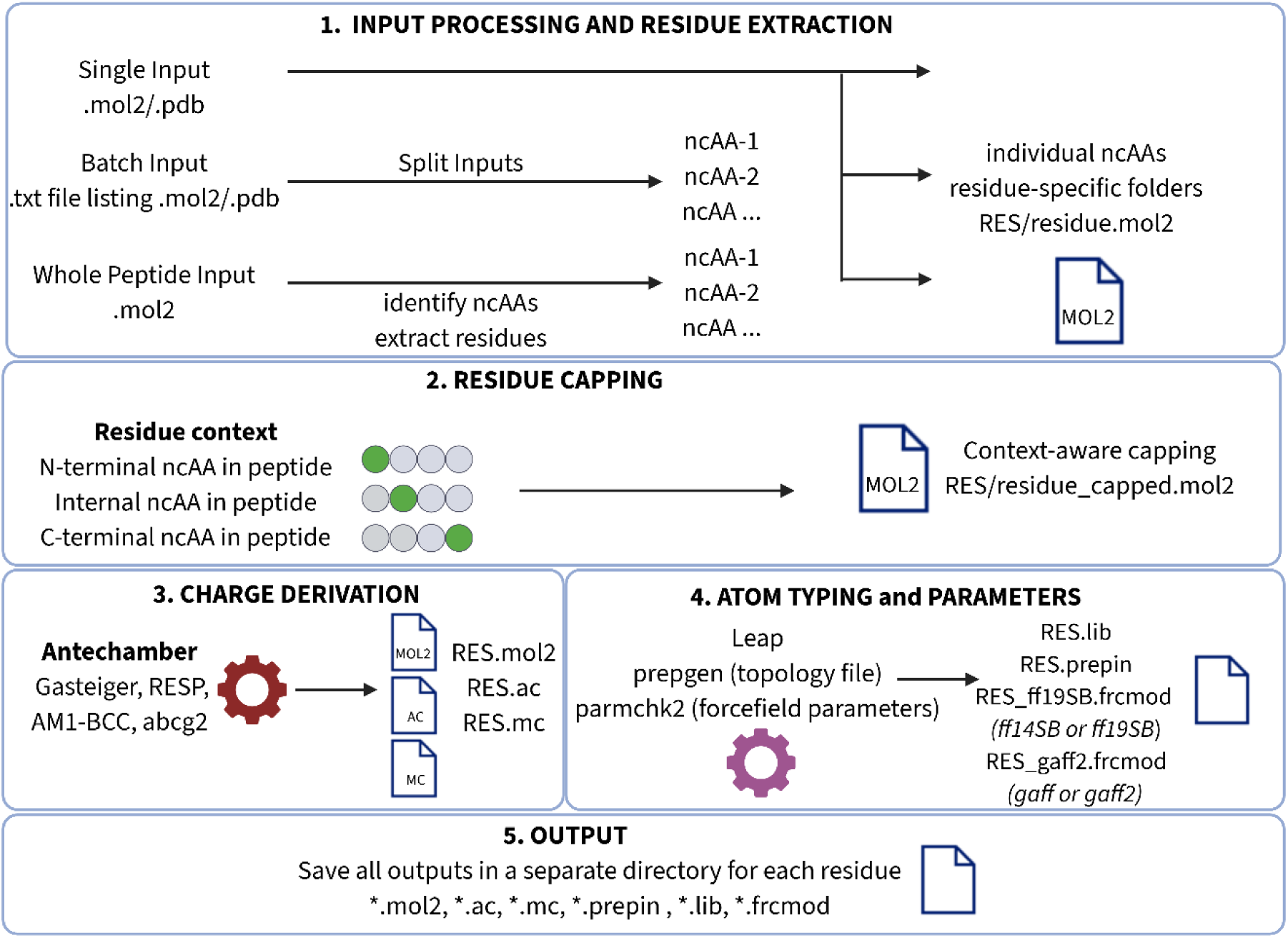
Overview of the TopFor workflow for automated parameterization of noncanonical amino acids. The workflow supports single-residue, whole-peptide, and batch input modes and performs residue isolation, terminal-context-aware capping, charge derivation, and atom typing, force-field parameter generation, and export of AMBER-compatible residue topology and parameter files for molecular dynamics simulations. Created with BioRender.

### Automated Parameterization Pipeline

The automated parameterization workflow (Figure 1) consists of four main stages: (1) input processing and residue extraction, (2) residue capping according to its position within the peptide, (3) charge derivation, and (4) atom typing and generation of residue templates and force-field parameter files. The implementation of each stage is described in the following sections.

TopFor is implemented as an open-source Python package and is available through GitHub (https://github.com/kalininalab/TopFor). The software is executed through a single command-line interface, with the input mode and charge-derivation method specified by the user. Software requirements, command-line usage, and output-file organization are described in the Supporting Information (Supplementary Note), with detailed installation instructions and example workflows provided in the GitHub repository.

### Input Processing and Residue Extraction

TopFor uses an individual ncAA residue as the basic unit for parameterization and supports three input modes: single-residue, batch, and whole-peptide processing. In single-residue mode, an individual ncAA is provided as a MOL2 or PDB file. PDB inputs are converted to MOL2 before parameterization, and each residue is processed independently in a dedicated output directory. By default, the input is treated as an internal residue, with its backbone nitrogen (head atom) and carbonyl carbon (tail atom) representing the two sites through which the residue forms peptide bonds with neighboring residues. The atom connecting the head and tail defines the main chain; for an α-amino acid, the Cα atom constitutes the main-chain atom between the backbone N and carbonyl C. If the input represents an *N*-terminal or *C*-terminal residue, the corresponding terminal context is specified by the user using the terminal-mode option.

Batch mode applies the same residue-level workflow independently to multiple input structures. The batch input may contain multiple MOL2 and/or PDB residue structures, which are processed in separate residue-specific output directories. Each input is therefore handled using the same procedure as in single-residue mode. Processing status is recorded independently for each residue, and failure of one residue does not interrupt processing of the remaining inputs. As in single-residue mode, residues are treated as internal by default, whereas *N*- or *C*-terminal context is specified by the user when required.

In whole-peptide mode, TopFor reads the atom, bond, and residue information from a complete peptide structure and identifies residues whose names are not included in the set of canonical amino acids. In contrast to single-residue and batch modes, the head, tail, main-chain, and terminal context of each ncAA are determined automatically from the peptide connectivity. TopFor identifies the residue atoms involved in peptide-bond connections to neighboring residues. For conventional α-amino acids, these correspond to the backbone N (head atom) and carbonyl C (tail atom). The main-chain atoms are then determined from the connectivity path between the head and tail atoms within the residue. An ncAA connected to neighboring residues through both its head and tail atoms is treated as an internal residue. The absence of a peptide-bond connection through the head identifies an *N*-terminal residue, whereas the absence of a connection through the tail identifies a *C*-terminal residue. For *C*-terminal residues, the presence of an OXT atom (the additional terminal carboxylate oxygen) is additionally detected and recorded for the capping step. Each identified ncAA is extracted as an individual MOL2 structure while retaining its coordinates and bonds between atoms within the residue. The automatically determined head, tail, main-chain, and terminal information is retained during extraction and used to select the appropriate capping procedure. Each extracted ncAA is then processed independently through the same residue-level parameterization workflow used in single-residue mode.

The current input-processing procedure was evaluated for ncAAs in linear, head-to-tail cyclic, and cyclic depsipeptide systems. In these systems, ncAAs connected through conventional peptide bonds can be processed automatically. Residues directly participating in nonstandard covalent linkages, such as the ester linkage in cyclic depsipeptides, currently require manual parameterization and definition of the corresponding linkage during system assembly, because TopFor does not automatically recognize these additional connections or determine the corresponding capping scheme.

### Residue Capping

Following residue extraction, TopFor generates a chemically complete capped structure for charge derivation and force-field parameter generation. Extraction of a residue from a peptide breaks the peptide backbone and leaves incomplete valences at the backbone nitrogen and carbonyl carbon. To reproduce the local chemical environment of a residue when incorporated into a peptide, TopFor adds appropriate capping groups while preserving the original Cartesian coordinates of the residue.

Capping is performed using a PyMOL-based procedure^43^. Existing hydrogen atoms are first removed from the residue structure. For internal residues, an acetyl group (ACE; CH₃CO–) is attached to the head atom, and an *N-*methyl group (NME; –NHCH₃) is attached to the tail atom. These caps mimic the local backbone environment of neighboring residues during charge derivation and parameter generation. For *N*-terminal residues, ACE capping is omitted to preserve the free amino terminus, whereas the tail atom is capped with NME. For C-terminal residues, the OXT atom is preserved and NME capping is omitted, whereas the head atom is capped with ACE. In whole-peptide mode, the appropriate capping scheme is selected automatically from the residue connectivity and terminal context determined during residue isolation. In single-residue and batch modes, the terminal context is specified by the user.

Following addition of the appropriate capping groups, hydrogen atoms are added using PyMOL^43^. TopFor evaluates the electron count of the capped structure against the net charge of the input residue. For negatively charged residues, a hydrogen is removed from a carboxyl oxygen using PyMOL, whereas for positively charged residues, a hydrogen is added to a suitable sidechain nitrogen using PyMOL. The residue net charge can also be specified explicitly by the user and is retained during processing. The resulting capped residue is then used for charge derivation and force-field parameter generation.

### Charge Derivation

TopFor supports four user-selectable charge derivation schemes: AM1-BCC^37,38^, ABCG2^39,40^, RESP^41^, and Gasteiger^42^. The selected charge method is applied to the capped residue structure generated in the preceding step. For AM1-BCC, ABCG2, and Gasteiger charges, TopFor calls the Antechamber module of AmberTools with the corresponding charge model and the net charge determined for the residue^35,44^.

AM1-BCC is used as the default charge model. ABCG2 and Gasteiger can be selected as alternative charge-derivation methods, with Gasteiger providing an empirical option that does not require quantum-mechanical charge calculations. For RESP charge derivation, TopFor uses a separate automated workflow that requires ORCA^45^ and Multiwfn^46^ to be installed separately by the user, while xTB^47^ is installed automatically as part of the TopFor Conda environment. The capped residue is first converted to XYZ format and subjected to an initial geometry optimization with xTB. The resulting structure is optimized with ORCA at the HF/6-31G* level, followed by a single-point calculation at the same level. The resulting wavefunction is converted to Molden^48^ format and used for RESP fitting with Multiwfn. The fitted charges are then transferred to the capped MOL2 structure using Antechamber. Because RESP requires additional geometry optimization, quantum-mechanical calculations, and charge fitting, its parameterization is computationally more demanding than the other supported charge derivation schemes.

### Atom Typing, Parameter Assignment and File Generation

Following charge derivation, TopFor generates the residue topology and force-field parameter files using Antechamber, prepgen, parmchk2, and LEaP from AmberTools^35^. Antechamber converts the charged MOL2 structure to AC format and assigns AMBER atom types. The resulting AC file contains the atom identities, atom types, connectivity, and partial charges required for residue-template generation.

TopFor then automatically prepares the main-chain definition (.mc) file using its internal prepgen_writer module. The .mc file is constructed from the residue topology determined during input processing and the capping information generated during residue preparation. It specifies the head and tail connection atoms, the main-chain atoms, the net residue charge, and the ACE and NME atoms to be omitted from the final residue template. The AC and .mc files are supplied to prepgen, which removes the specified capping atoms and generates the amino-acid residue template (.prepin).

TopFor employs parmchk2 to identify force-field parameters required for the ncAA. The residue is evaluated separately against a user-selected protein force-field parameter database and a user-selected general force-field parameter database. TopFor currently supports ff99SB^49^, ff14SB^50^ and ff19SB^51^ as protein force fields and GAFF^37,38^ and GAFF2^39^ as general force fields, with ff19SB and GAFF2 used by default. For each selected force field, parmchk2 compares the bond, angle, dihedral, improper, and nonbonded parameters required by the residue with those available in the corresponding parameter database and writes parameters to a force-field modification (.frcmod) file. Problematic parameters that cannot be adequately assigned are marked with “*ATTN, need revision*” and require inspection before use.

TopFor generates both protein- and general-force-field (.frcmod) files because the selected protein force field may not contain all parameters required for the noncanonical chemical features of an ncAA. The resulting files allow the user to inspect the available parameterization and select the protein-force-field (.frcmod), the general-force-field (.frcmod), or both files as appropriate for system preparation.

LEaP is used within TopFor to load the final MOL2 structure together with the generated protein- and general-force-field .frcmod files and to generate a library file (.lib) and test topology and coordinate files (.prmtop and .inpcrd). This step provides a residue-level compatibility check within the AMBER framework.

For each successfully processed residue, TopFor retains the intermediate and final files generated during parameterization, including the charged MOL2 and AC structures, main-chain definition (.mc), residue template (.prepin), protein and general-force-field .frcmod files, residue library (.lib), and test topology and coordinate files (.prmtop and .inpcrd). A residue-specific metadata file (residue_meta.json) additionally records the residue topology, terminal context, applied capping strategy, net charge, selected charge model, and validation information. Representative intermediate and final files generated during the parameterization of *N-*methyl-L-valine (MVA) are provided in Tables S8-15 to illustrate the file transformations and AMBER-compatible outputs produced by the workflow.

### Validation and Application Protocols

Validation was performed using linear peptides composed of canonical amino acids as a baseline scenario, followed by evaluation of TopFor with head-to-tail cyclic and cyclic depsipeptide systems and in high-throughput peptide-design workflows. The systems used to evaluate TopFor, together with their peptide architecture, parameterization mode, charge-derivation method, simulation protocol, and purpose, are summarized in Table S3.

### Validation Using Canonical Amino Acids in a Linear Peptide: MDM2-p53 complex

To evaluate whether residue definitions generated through the automated TopFor workflow are compatible with AMBER simulations and preserve the structural behavior observed with native AMBER residue definitions, canonical amino acids were used as reference residues. Because these residues have definitions in the AMBER force field, they provide a direct reference against which residues independently parameterized through TopFor can be evaluated. Selected canonical residues were therefore renamed while their atoms, bonds, and coordinates were left unchanged (e.g., THR was renamed THX). The renamed residues were no longer recognized as canonical amino acids by TopFor and were processed as noncanonical residues through the complete parameterization workflow. This strategy allowed chemically identical residues represented either by the native AMBER definitions or by residue definitions prepared through TopFor to be compared directly, thereby isolating the effect of the automated parameterization procedure from differences in residue chemistry.

The experimentally determined complex of MDM2 with a linear p53 peptide (PDB ID: 1YCR^52^) was processed using the whole-peptide input mode of TopFor. For residue types occurring multiple times within the peptide, the same parameterized residue definition was reused for each occurrence. Four variants were compared. The STANDARD variant retained the native AMBER residue definitions for the p53 peptide, whereas the other variants used residue templates and parameter files prepared through the automated TopFor workflow using AM1-BCC, ABCG2, or RESP partial charges. MD simulations were performed as described in Section “Molecular Dynamics Simulations”. Parameterization timings were measured on a workstation equipped with an Intel Xeon w7-3455 processor and 125 GiB of system memory.

### Application to the Head-to-Tail Cyclic Peptide: Cyclophilin–Cyclosporin A Complex

To evaluate the applicability of TopFor to head-to-tail cyclic peptides, the crystal structure of the cyclophilin–cyclosporin A complex from *Trypanosoma cruzi* (PDB ID: 1XQ7^53^) was selected. Cyclosporin A is an 11-residue head-to-tail cyclic peptide containing nine noncanonical residue positions comprising six unique ncAA types: L-α-aminobutyric acid (ABA), 4-methyl-4-[(E)-2-butenyl]-4,*N*-methylthreonine (BMT), D-alanine (DAL), *N*-methylleucine (MLE), MVA, and sarcosine (SAR).

The ncAAs of cyclosporin A were processed using the batch input mode of TopFor with ABCG2 settings. One representative of each unique residue type was parameterized through the automated workflow, and the resulting residue templates and parameter files were reused for repeated residue types within the peptide. The parameterized residues were incorporated into the head-to-tail cyclic peptide during system preparation in LEaP. MD simulations of the resulting cyclophilin–cyclosporin A complex were performed as described in Section “Molecular Dynamics Simulations”.

### Application to a Cyclic Depsipeptide: DnaN-Cyclohexylgriselimycin Complex

To evaluate the applicability of TopFor to cyclic depsipeptides, the crystal structure of the DnaN–CGM complex (PDB ID: 8CIY^15^) was selected. CGM is a cyclic depsipeptide containing seven ncAA residues representing five ncAA types: *N*-methyl-D-leucine (MLU), (4R)-4-methyl-L-proline (MP8), MVA, *N*-methylidene-L-threonine (NZC), and (4S)-4-cyclohexyl-L-proline (PH6). Macrocycle formation involves an ester linkage between NZC and GLY. The ncAAs connected through conventional peptide bonds were parameterized individually using the single-residue input mode of TopFor. In contrast, the two residues directly involved in the depsipeptide ester linkage, NZC and GLY, were parameterized manually because this sidechain-to-backbone connectivity is not currently supported by the TopFor automated workflow. For these residues, the required capping structures and main-chain (.mc) definitions were prepared manually to represent the ester-linked connectivity, and parameterization was performed directly using AmberTools^35^ utilities, including Antechamber, prepgen, and parmchk2, without TopFor. The resulting residue templates (.prepin) and force-field modification files (.frcmod), together with TopFor-generated parameters for the remaining ncAAs, were used to prepare the complete DnaN–CGM system. MD simulations were performed as described in Section “Molecular Dynamics Simulations”.

### Integration of TopFor into High-Throughput Virtual Screening of DnaN–Cyclohexylgriselimycin Derivatives

To demonstrate the integration of TopFor into a high-throughput computational peptide discovery workflow, virtual screening was performed using the DnaN–CGM complex (PDB ID: 8CIY^15^). Libraries of CGM derivatives were generated by modifying selected positions of the native peptide scaffold, docked into DnaN using Glide^54^, and parameterized through the automated TopFor workflow. The resulting complexes were subjected to AMBER^36^-based post-docking refinement and single-frame Molecular Mechanics/Generalized Born Surface Area (MM/GBSA) evaluation, followed by short MD simulations of the highest-ranked candidates.

The DnaN structure was prepared using the Protein Preparation Wizard in Schrödinger Maestro^55^. A docking grid of 20 × 20 × 20 Å was centered on the native CGM binding site. Commercially available α-amino acid and carboxylic acid building blocks were obtained from the Enamine^56^ catalogue and used to construct the corresponding CGM derivative libraries. The generated structures were prepared in MOE^57^ by structure washing, partial-charge assignment, and energy minimization. Glide docking was performed using a core-constrained protocol in which the conserved CGM scaffold was defined by a SMARTS pattern, while structural variation was permitted at the derivatized position. As docking was used primarily to generate initial three-dimensional protein–peptide complex structures for AMBER^36^-based refinement, only the highest-ranked pose was retained for each derivative and Glide post-docking minimization was disabled.

Two independent derivatization campaigns were performed. The first investigated substitution of the MVA1 residue using 3,194 commercially available α-amino acid building blocks from the Enamine^56^ catalogue. Following physicochemical filtering and Glide docking, 521 candidate derivatives were parameterized using the batch input mode of TopFor. The second campaign investigated derivatization of the acetyl group using 75,132 commercially available carboxylic acid building blocks from the Enamine^56^ catalogue. Following physicochemical filtering and Glide docking, 693 candidate derivatives were parameterized using the same workflow. For both campaigns, the manually prepared N-methylthreonine (NZC) and custom glycine (GLZ) residue templates and their corresponding force-field modification files were used during LEaP assembly to retain the depsipeptide ester linkage. Glycine was renamed from GLY to GLZ to define it as a distinct residue with the custom connectivity required for ester-bond formation rather than the peptide-bond connectivity of the standard AMBER glycine template.

Following parameterization and system assembly, the docked complexes were subjected to a two-stage AMBER minimization to refine the initial docking geometries before MM/GBSA evaluation using MMPBSA.py^58^. Each complex was first subjected to 3,000 minimization cycles, with the first 1,000 cycles performed using steepest descent followed by conjugate-gradient minimization, while a positional restraint of 10 kcal mol⁻¹ Å⁻² was applied to the protein. A 10 Å nonbonded cutoff was used. This was followed by an unrestrained minimization of 4,000 cycles, comprising 2,000 cycles of steepest descent followed by conjugate-gradient minimization with the same 10 Å cutoff. Both minimization stages were performed using pmemd.cuda. Single-frame MM/GBSA calculations were performed on the final minimized structures using MMPBSA.py with the GBneck2 implicit-solvent model (igb=8)^58^. For this high-throughput application, a derivative was considered successfully processed when TopFor parameterization was completed and the resulting DnaN–CGM complex completed the two-stage AMBER post-docking minimization and single-frame MM/GBSA calculation. The workflow completion rates reported for the two derivatization campaigns were determined using this criterion.

Based on the resulting MM/GBSA ranking, the top 50 peptide derivatives from each library were selected for 5 ns production MD simulations performed according to the protocol described in the Molecular Dynamics Simulations section. Following MD, MM/GBSA calculations were performed over the final 2 ns of each 5 ns trajectory using MMPBSA.py^58^ with the GBneck2 implicit-solvent model (igb=8). These short 5 ns simulations were used to demonstrate the applicability of TopFor-generated residue parameterizations in a high-throughput virtual screening workflow, including AMBER equilibration, production MD, and trajectory-based MM/GBSA analysis, rather than for comprehensive lead optimization.

Chemical-space coverage of the successfully processed derivatives was evaluated using Morgan fingerprints generated with RDKit^59^, with a radius of 2, a fingerprint length of 2,048 bits, and inclusion of chirality. The resulting binary fingerprints were projected into two dimensions using Uniform Manifold Approximation and Projection (UMAP)^60^ with the Jaccard distance metric, 20 nearest neighbors, a minimum distance of 0.15, and a random seed of 42.

### Structure-Based Design and Computational Evaluation of GM-Derived Peptides

The crystal structure of *Escherichia coli (E. coli)* DnaN in complex with griselimycin (GM; PDB ID: 8CIX^15^) was used as the starting structure. Noncanonical amino acids within GM were parameterized using TopFor, and the DnaN–GM complex was prepared according to the protocol described in the Molecular Dynamics Simulations section. The complex was subjected to a 1000 ns production MD simulation. Trajectory analysis was performed using cpptraj^61^. MM/GBSA per-residue energy decomposition was performed over the final 200 ns (800–1000 ns) of the production trajectory using MMPBSA.py^58^ with the GBneck2 implicit-solvent model (igb=8).

A focused library of 89 GM-derived peptide candidates was constructed, comprising 61 tetrapeptides, 21 pentapeptides, and seven N-terminally fatty-acid-modified derivatives. The generated compounds were docked into the GM-binding site of *E. coli* DnaN using the Standard Precision (SP) protocol of Glide (Schrödinger)^54^. The receptor grid was generated using the Receptor Grid Generation tool in Schrödinger^54^ and defined as a 20 × 20 × 20 Å box centered on the crystallographic GM ligand. The resulting protein–peptide complexes were prepared for AMBER^36^ simulations using the parameterization and system-preparation procedures described in the Molecular Dynamics Simulations section. Each complex was subjected to the two-stage energy minimization, heating, density equilibration, equilibration, and 10 ns production MD protocol described therein.

MM/GBSA binding-energy calculations were performed using MMPBSA.py^58^ with the GBneck2 implicit-solvent model (igb=8) at successive stages of the computational workflow. Single-frame calculations were performed using the structures obtained after the first (emin1) and second (emin2) energy-minimization stages. During the 10 ns production trajectories, MM/GBSA calculations were performed for frames 1–250, 251–499, and 500–999, with every tenth frame included in each calculation. An additional single-frame MM/GBSA calculation was performed using the final structure of each production trajectory. The calculated MM/GBSA energies together with structural inspection of the protein–peptide complexes were used for compound selection.

Following experimental evaluation against *M. smegmatis* DnaN by SPR, EFB1–EFB6 were docked into *M. smegmatis* DnaN (PDB ID: 5AH2) to examine their predicted binding modes in the experimentally evaluated target. Docking was performed using the Glide SP protocol. A 20 × 20 × 20 Å receptor grid was generated using the Receptor Grid Generation tool and centered on the crystallographic GM ligand. Because EFB1–EFB6 retain the ACE–MVA–Pro motif of GM, the common ACE–MVA–Pro substructure was defined using a SMARTS pattern and constrained to the corresponding region of the crystallographic GM binding pose during docking. The resulting binding poses were visually inspected to examine peptide orientation within the DnaN binding groove and interactions with neighboring residues.

### Molecular Dynamics Simulations

All MD simulations were performed using the AMBER24 simulation package^36^. Systems were prepared using the ff19SB^51^ protein force field, GAFF2^39^, and the OPC^62^ explicit water model. After system neutralization, each system underwent a two-stage energy minimization. In the first stage, 3,000 minimization cycles were performed, comprising 1,000 cycles of steepest descent followed by 2,000 cycles of conjugate-gradient minimization, with a positional restraint of 10 kcal mol⁻¹ Å⁻² applied to the protein. This was followed by an unrestrained minimization of 4,000 cycles, comprising 2,000 cycles of steepest descent followed by 2,000 cycles of conjugate-gradient minimization. A 10 Å nonbonded cutoff was applied during both minimization stages.

The minimized systems were heated from 0 to 300 K over 50 ps under constant-volume conditions with a positional restraint of 2 kcal mol⁻¹ Å⁻² applied to the protein. This was followed by 100 ps of constant-pressure density equilibration at 1 bar with the restraint reduced to 1 kcal mol⁻¹ Å⁻² and a further 200 ps of constant-pressure equilibration with a 0.5 kcal mol⁻¹ Å⁻² restraint. Pressure was controlled using the Berendsen barostat^63^ with isotropic position scaling and a pressure relaxation time of 5 ps. The restraints were removed for the production simulations. These settings correspond to the heating, density-equilibration, and equilibration stages specified in the simulation protocol.

Production simulations were performed under constant-pressure conditions at 300 K and 1 bar using the same pressure-control settings. A 2 fs integration time step was used, with bonds involving hydrogen constrained using the SHAKE^64^ algorithm. Long-range electrostatic interactions were treated using the particle mesh Ewald (PME)^65,66^ method with a 10 Å real-space cutoff.

Temperature was maintained using a Langevin thermostat^67^ with a collision frequency of 2 ps⁻¹. Production simulations were performed using the GPU-accelerated pmemd.cuda engine, and coordinates were written every 10 ps for trajectory analysis.

The production simulation length depended on the application. The MDM2–p53 (PDB 1YCR^52^) validation systems were simulated for 100 ns in three independent replicates, whereas the cyclophilin– cyclosporin A (PDB 1XQ7^53^) and DnaN–CGM (PDB 8CIY^15^) complexes were each simulated for a single 100 ns trajectory. For the high-throughput virtual screening workflow, the two-stage minimization described above had already been performed as part of the post-docking refinement and MM/GBSA ranking procedure and was therefore not repeated for the selected compounds. Instead, the minimized structures of the top 50 derivatives from each derivatization campaign were continued from the heating stage through density equilibration and equilibration, followed by 5 ns of production MD using the protocol described above. The *E. coli* DnaN–GM complex (PDB ID: 8CIX^15^) was subjected to a single 1000 ns production simulation for the structure-based peptide discovery application, whereas each of the 89 GM-derived peptide candidates was subjected to a 10 ns production simulation.

Trajectory analyses were performed using cpptraj^61^ and included backbone root-mean-square deviation (RMSD), root-mean-square fluctuation (RMSF), radius of gyration (Rg), backbone Ramachandran (ϕ/ψ) distributions, and secondary-structure analysis, together with visual inspection of peptide conformations.

Representative structures were obtained by k-means clustering using cpptraj. Prior to clustering, trajectories were autoimaged, solvent and counterions were removed, and frames were aligned to the receptor backbone using the Cα, C, and N atoms. Clustering was subsequently performed on the heavy atoms of the bound peptide using RMSD as the distance metric without additional fitting. Ten clusters were generated using a random sieve interval of 10 frames. For the MDM2–p53, cyclophilin– cyclosporin A, and DnaN–CGM systems, clustering was performed over the final 50 ns of the 100 ns production trajectories. For the 1000 ns *E. coli* DnaN–GM simulation, clustering was performed over the complete production trajectory. For structural comparison with the corresponding experimental crystal structure, the representative frame of the most populated cluster was used.

Compound-level results from the high-throughput workflow, including UMAP coordinates, post-docking minimization and MM/GBSA data, and trajectory-based MM/GBSA values for the selected derivatives, are provided in Supplementary Data 1.

### Peptide Synthesis

EFB1–EFB6 were custom-synthesized by GenScript and supplied at ≥90% purity, with 14 mg of each peptide provided. Peptide identity and purity were verified by the manufacturer as part of the quality-control procedure. The peptide sequences are provided in Table S7.

### Surface Plasmon Resonance (SPR) Measurements

Surface plasmon resonance (SPR) measurements were performed using a Biacore X100 system (GE Healthcare). *M. smegmatis* DnaN was diluted to 100 µg/mL in sodium acetate buffer (pH 4.5) and immobilized on a CM7 sensor chip by amine coupling, resulting in a final immobilization level of approximately 13,970 response units (RU).

For the initial binding screen, EFB1–EFB6 were tested at 500 µM in HBS-EP buffer containing 1% DMSO. GM was additionally measured at 100 nM as a reference compound during the initial exploratory measurement. SPR responses were determined after reference subtraction (Fc2−Fc1). EFB4, EFB5, and EFB6, which showed the strongest responses in the initial screen, were then evaluated over a concentration range of 62.5 µM – 1 mM. Samples were injected at a flow rate of 30 µL/min with a contact time of 180 s, followed by a dissociation time of 600 s. Reference-subtracted sensorgrams (Fc2−Fc1) were used for analysis. As the measurements did not permit reliable determination of equilibrium dissociation constants (*K*D), the SPR data were interpreted qualitatively based on concentration-dependent SPR responses rather than as quantitative affinity measurements.

## Results

### Residues parameterized through TopFor preserve the structural and conformational behavior of the linear MDM2–p53 complex

The MDM2–p53 complex (PDB ID: 1YCR)^52^ was used to assess whether automation of residue parameterization through TopFor affects the structural behavior obtained with the native AMBER representation. Selected canonical residues of the bound p53 peptide were renamed without altering their chemical structures and processed through TopFor as ncAAs, providing a direct comparison with their native AMBER definitions. AM1-BCC, ABCG2, and RESP parameterizations were compared with the native AMBER representation (STANDARD), with three independent 100 ns production trajectories performed for each system.

Representative structures obtained by clustering the equilibrated portions of each independent trajectory were superposed with the experimental crystal structure after alignment on the MDM2 backbone (Figure 2). Across the representative structures, MDM2 backbone RMSDs relative to the crystal structure ranged from 1.06 to 1.80 Å, while p53 backbone RMSDs calculated after MDM2 backbone fitting ranged from 1.22 to 1.73 Å (Table S1). The superpositions retained the α-helical conformation of the bound p53 peptide and the orientations of the hydrophobic anchor residues across all four parameterization schemes, with no systematic parameterization-dependent structural rearrangements.

**Figure 2.**
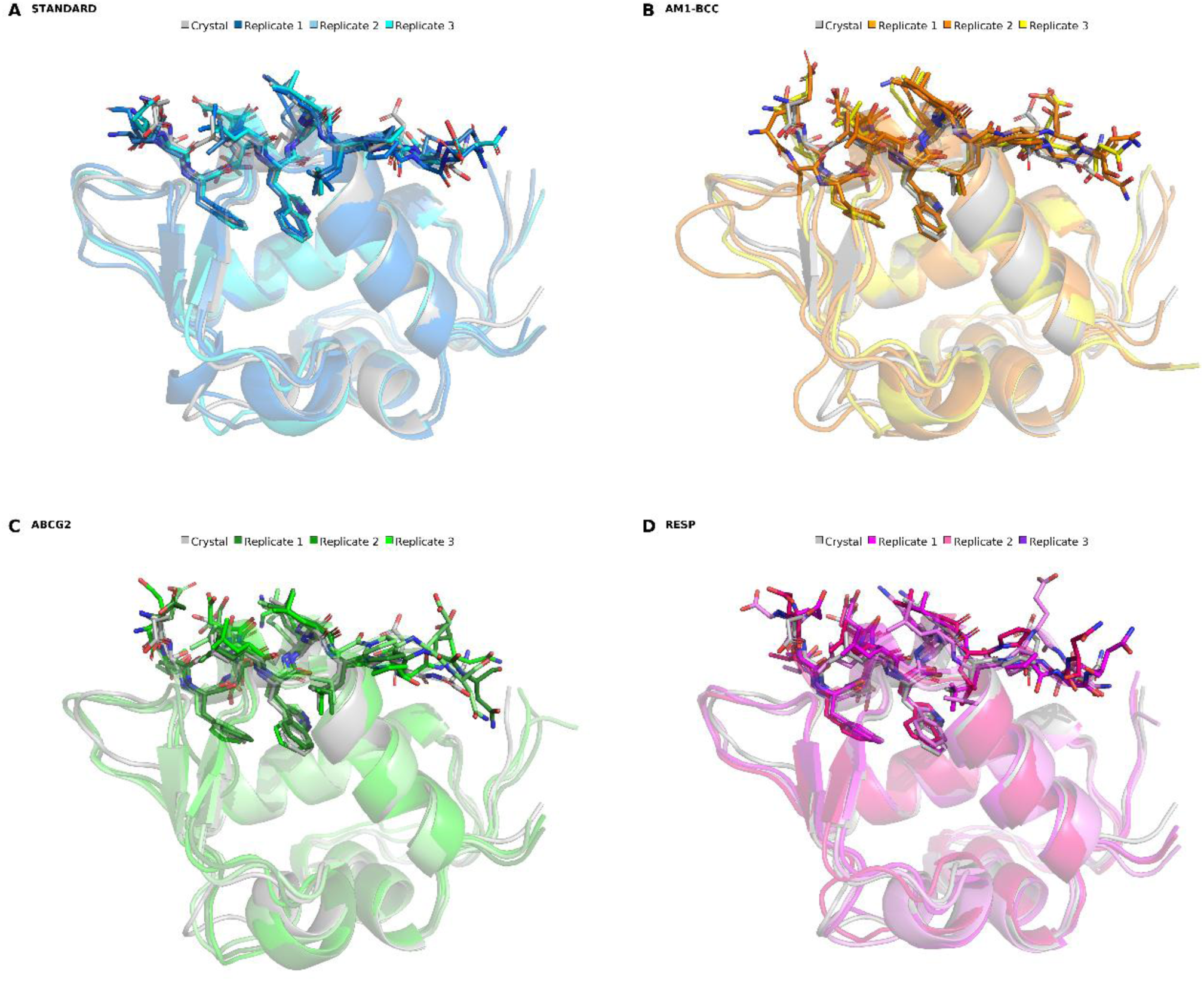
Validation of TopFor-generated parameterizations using the canonical linear MDM2–p53 complex (PDB ID: 1YCR). Representative structures obtained by clustering the final 50 ns of each of the three independent 100 ns production trajectories are shown for the (A) STANDARD, (B) AM1-BCC, (C) ABCG2, and (D) RESP parameterization schemes. The experimental crystal structure is shown in gray, while representative structures from the three independent simulations are shown in different shades corresponding to each parameterization method. Structures were aligned using the MDM2 backbone. MDM2 is shown as a cartoon representation and the bound p53 peptide as sticks. The superposition illustrates preservation of the experimentally observed peptide-binding mode and overall structural organization of the MDM2–p53 complex across the four parameterization schemes.

All twelve simulations completed without numerical instability. Mean MDM2 backbone RMSDs ranged from approximately 1.1 to 1.4 Å across the individual trajectories, while p53 peptide RMSDs ranged from approximately 0.9 to 1.1 Å (Figure S1). Mean complex radii of gyration ranged from 13.06 to 13.15 Å, with within-trajectory standard deviations of 0.06–0.09 Å (Figure S3).

Backbone RMSF profiles were consistent across the four parameterization schemes, with the p53 hydrophobic anchor residues Phe88 and Trp92 showing backbone RMSF values below 2 Å across all parameterization schemes and replicates (Figure S2). The four schemes populated the same dominant regions of φ/ψ space (Figure S4). DSSP^61^ analysis showed >80% α-helical occupancy for Phe3 and nearly 100% for Ser4–Trp7, while the remaining residues were annotated as coil (Figure S5). The TopFor-parameterized systems therefore showed structural and conformational behavior comparable to the native AMBER reference over the simulated timescale.

In our calculations, parameterization of the *N*-methylvaline (MVA) test residue required approximately 2 s with Gasteiger charges, 10 s with either AM1-BCC or ABCG2, and 15 min with RESP. The lower computational cost of AM1-BCC and ABCG2 supported their use for more complex ncAA-containing systems and large-scale virtual screening.

### TopFor enables parameterization of the head-to-tail cyclic cyclophilin–cyclosporin A complex

Following validation with the linear MDM2–p53 system, we next examined whether TopFor could be applied to a structurally more complex head-to-tail cyclic peptide containing multiple ncAAs. The experimentally resolved *Trypanosoma cruzi* cyclophilin–cyclosporin A complex (PDB ID: 1XQ7^53^) was selected for this purpose. Cyclosporin A is an 11-residue head-to-tail cyclic peptide containing nine noncanonical residue positions (Figure 3C). The ncAAs were parameterized using the batch input mode of TopFor with ABCG2 partial charges.

**Figure 3.**
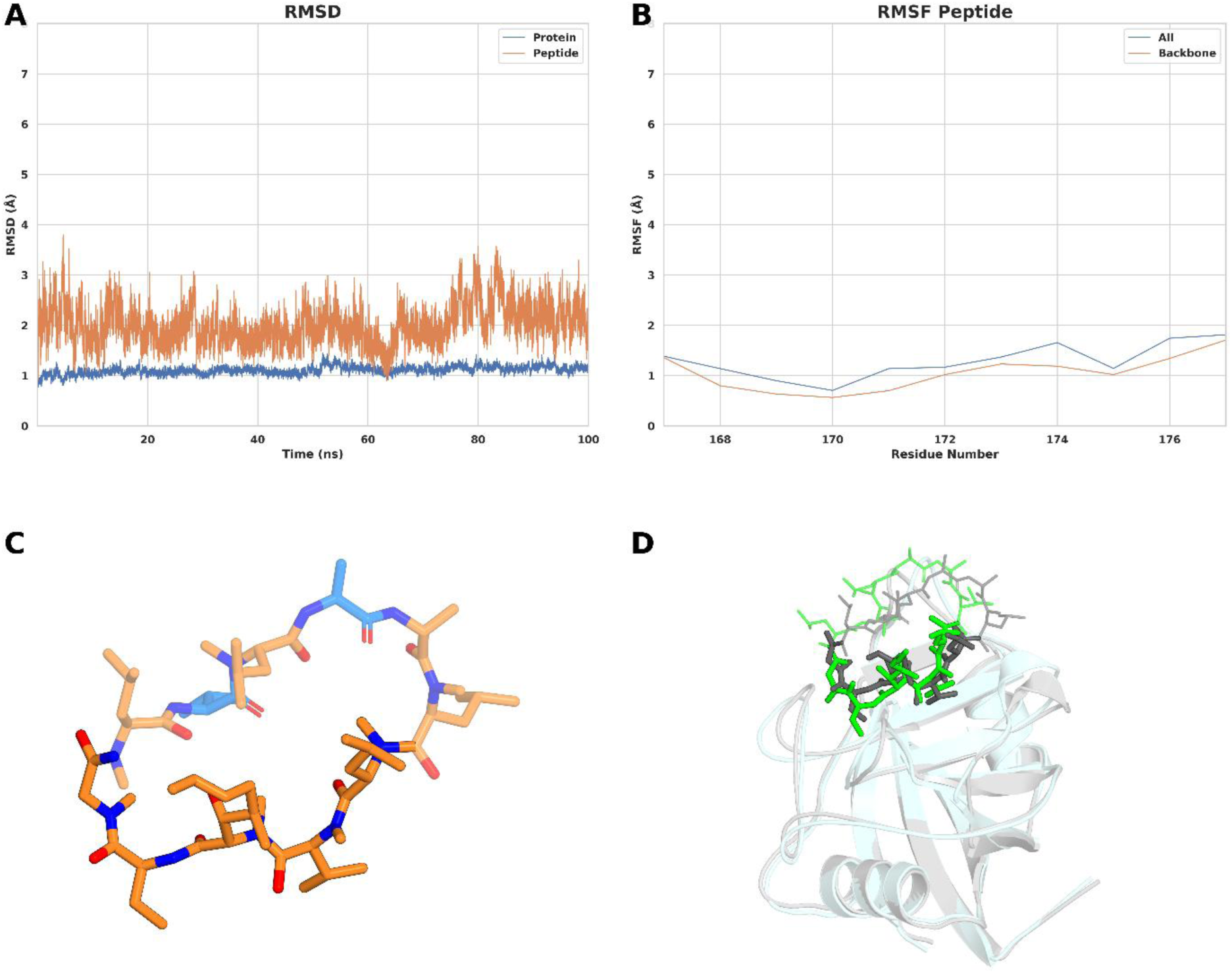
Application of TopFor to the cyclophilin–cyclosporin A complex (PDB ID: 1XQ7). (A) Backbone RMSD of cyclophilin and the bound cyclosporin A peptide during the 100 ns production simulation. (B) Backbone RMSF of the bound cyclosporin A peptide. (C) Structure of cyclosporin A showing the nine noncanonical amino acid positions parameterized using TopFor in orange and the two canonical amino acids in marine. Nitrogen and oxygen atoms are shown in blue and red, respectively. (D) Superposition of the representative structure obtained by clustering the final 50 ns of the production trajectory with the experimental crystal structure. Cyclophilin is shown as a cartoon, with the crystal structure in light gray and the MD representative structure in pale cyan. cyclosporin A is shown as sticks, with the crystal structure in gray and the MD representative structure in green. The cyclophilin-binding segment of cyclosporin A (residues 169–173) is emphasized with thicker sticks, whereas the solvent-exposed residues (167–168 and 174–177) are shown with thinner sticks.

The simulation completed without numerical instability. The cyclophilin backbone remained close to the starting structure throughout the 100 ns production trajectory, with a mean RMSD of 1.11 ± 0.08 Å (Figure 3A). Cyclosporin A showed greater deviation from the starting structure, with a mean RMSD of 2.01 ± 0.40 Å and transient excursions reaching 3.80 Å. Residue-level RMSF values remained below approximately 2 Å across the peptide (Figure 3B), and the radius of gyration showed no systematic change over the simulation (Figure S6). Superposition of the representative structure with the experimental crystal structure gave RMSDs of 0.55 Å for cyclophilin and 2.86 Å for the complete cyclosporin A peptide. The cyclophilin-binding segment of cyclosporin A (residues 169–173) showed an RMSD of 1.84 Å, whereas the solvent-exposed residues showed greater deviation (3.56 Å), indicating that the structural difference was concentrated primarily in the solvent-exposed region (Figure 3D). TopFor therefore enabled automated parameterization of multiple ncAAs within the head-to-tail cyclic peptide and MD simulation of the complete cyclic system.

### TopFor enables parameterization of Cyclohexylgriselimycin in the DnaN–CGM complex

We then evaluated TopFor using CGM, a cyclic depsipeptide containing multiple ncAAs and an ester linkage responsible for macrocycle formation. The experimentally resolved *E. coli* DnaN–CGM complex (PDB ID: 8CIY^15^) was selected for this purpose. Four ncAAs connected through conventional peptide bonds (MVA, MP8, MLU, and PH6) were parameterized through the TopFor single-residue workflow using AM1-BCC partial charges. The two residues directly involved in the depsipeptide ester linkage, NZC and Gly, were parameterized manually because this sidechain-to-backbone ester connectivity is not currently supported by TopFor (Figure 4C). Gly was renamed GLZ to define it as a distinct residue with the custom connectivity required for ester-bond formation.

**Figure 4.**
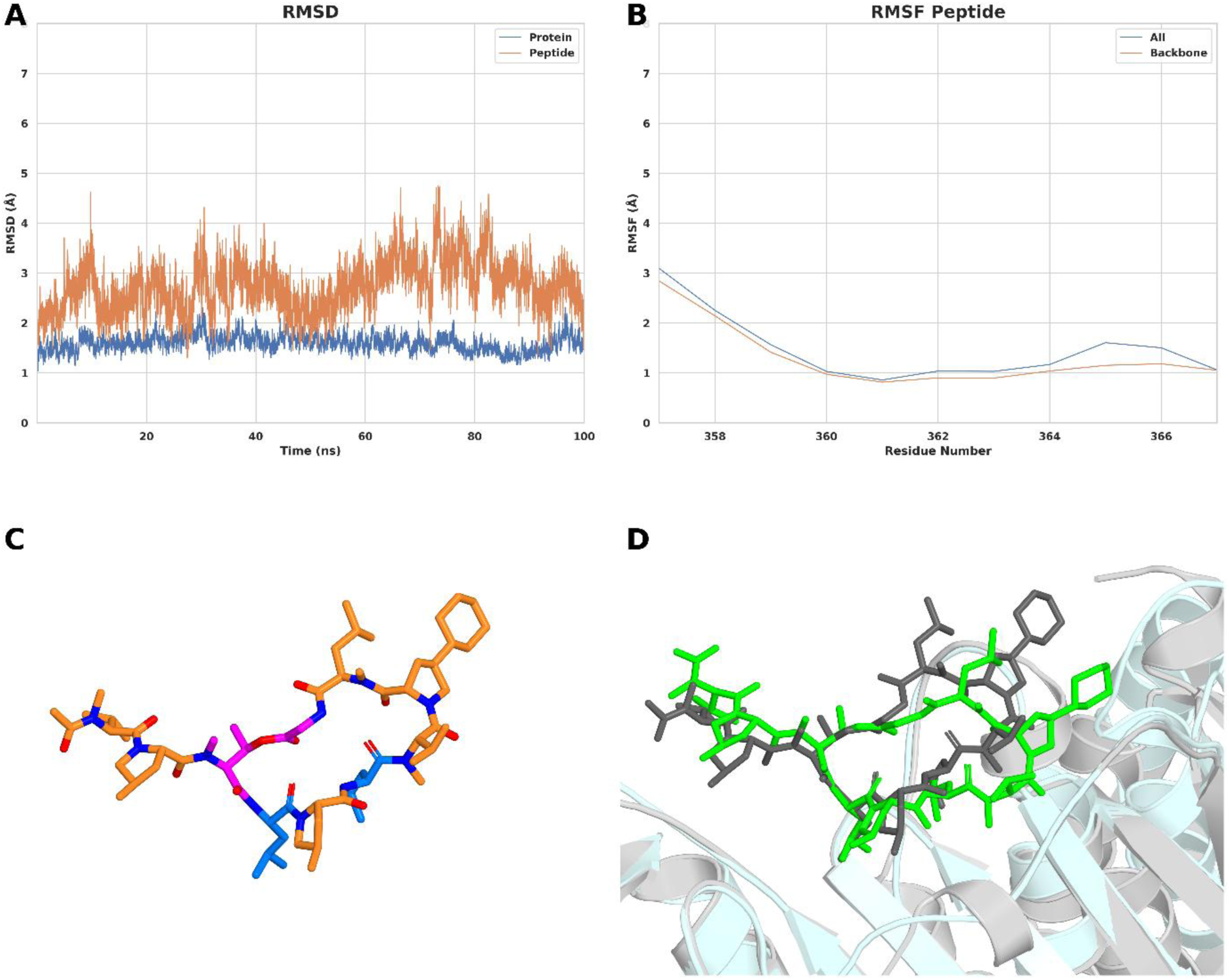
Application of TopFor to the CGM in the DnaN–CGM complex (PDB 8CIY). (A) Backbone RMSD of DnaN and the bound CGM peptide during the 100 ns production simulation. (B) Backbone RMSF of the bound CGM peptide. (C) Structure of CGM highlighting the noncanonical amino acids parameterized using TopFor in orange and the canonical amino acids in marine. The ester-linked residues requiring manual parameterization are highlighted in magenta. Nitrogen and oxygen atoms are shown in blue and red, respectively. (D) Superposition of the representative structure obtained by clustering the final 50 ns of the production trajectory with the experimental crystal structure. DnaN is shown as a cartoon, with the crystal structure in light gray and the MD representative structure in pale cyan. CGM is shown as sticks, with the crystal structure in gray and the MD representative structure in green.

During the 100 ns production trajectory, the DnaN backbone showed a mean RMSD of 1.60 ± 0.17 Å, whereas CGM showed a mean RMSD of 2.73 ± 0.50 Å, with transient excursions reaching 4.75 Å (Figure 4A). Residue-level RMSF analysis localized the largest CGM backbone fluctuations to the N-terminal region, comprising ACE (residue 357) and MVA (residue 358), with RMSF values of 2.84 and 2.14 Å, respectively; backbone RMSF values for the remaining residues ranged from 0.82 to 1.41 Å (Figure 4B). The radius of gyration showed no systematic change over the simulation (Figure S7). Superposition of the representative MD structure with the experimental crystal structure, following alignment on the DnaN backbone atoms, gave RMSDs of 1.49 Å for the DnaN backbone atoms and 2.36 Å for the CGM backbone (Figure 4D). Hence, TopFor can automate the parameterization of supported ncAAs within CGM and enable their incorporation into an AMBER MD workflow for the complete DnaN–CGM complex. Residues directly involved in sidechain-to-backbone ester-bond formation remain a current limitation because these additional covalent connections are not automatically recognized by TopFor and therefore require manual parameterization.

### TopFor enables high-throughput parameterization of CGM derivatives for downstream MD simulation workflows

Having established the applicability of TopFor to the DnaN–CGM system, we next examined its use for large-scale parameterization of CGM derivatives. Two derivatization sites representing different chemical environments were investigated: substitution of the internal MVA1 residue and modification of the N-terminal acetyl group (Figure 5A). AM1-BCC was used for charge derivation in both campaigns.

**Figure 5.**
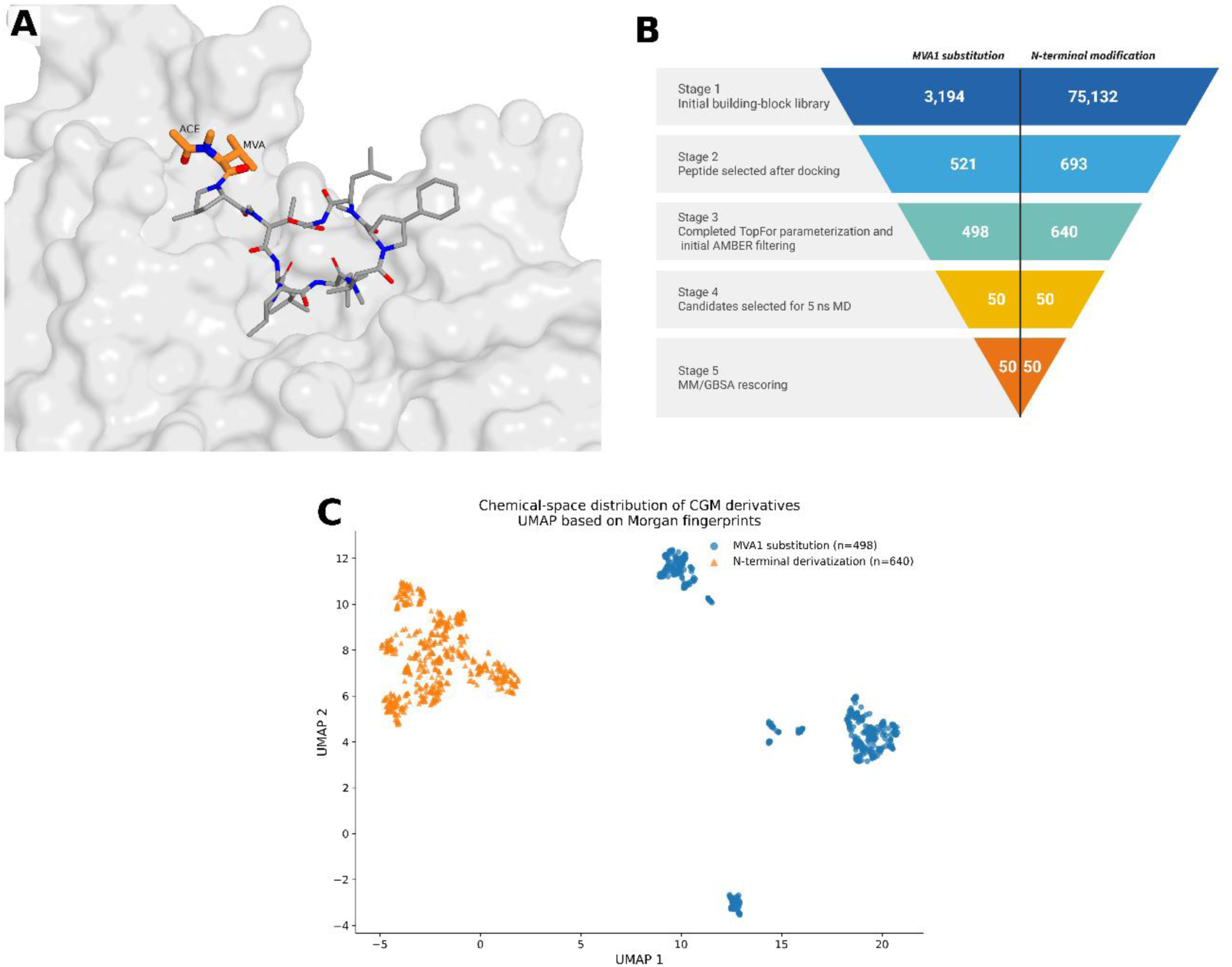
High-throughput application of TopFor to chemically diverse CGM derivatization libraries. (A) DnaN– CGM complex highlighting the two positions selected for chemical diversification: the MVA1 residue and the N-terminal acetyl (ACE) group. DnaN is shown as a gray surface representation and CGM as sticks, with MVA1 and ACE highlighted to indicate the derivatization sites. (B) Summary of the high-throughput virtual screening (HTVS) workflows for the MVA1 substitution and N-terminal (ACE) derivatization campaigns. Created with BioRender. (C) UMAP projection of derivatives that completed TopFor parameterization and the initial AMBER-based filtering workflow for the MVA1 substitution (*n* = 498) and N-terminal derivatization (*n* = 640) campaigns. Each point represents one CGM derivative and is colored according to the corresponding derivatization strategy. The projection illustrates the distribution of processed derivatives across the projected chemical space for the two derivatization strategies.

For MVA1 substitution, screening of derivatives generated from 3,194 commercially available α-amino acid building blocks by physicochemical filtering and core-constrained Glide^54^ docking identified 521 candidates for parameterization. Of these, 498 (95.6%) completed TopFor parameterization and the AMBER^36^-based post-docking refinement and single-frame MM/GBSA evaluation, whereas 23 (4.4%) did not complete the workflow. For *N*-terminal derivatization, screening of derivatives generated from 75,132 commercially available carboxylic acid building blocks using the same strategy identified 693 candidates. Of these, 640 (92.4%) completed TopFor parameterization and the AMBER^36^-based refinement and single-frame MM/GBSA evaluation, whereas 53 (7.6%) did not complete the workflow.

Overall, 1,138 of 1,214 derivatives (93.7%) completed TopFor parameterization and the initial AMBER^36^-based filtering workflow across the two campaigns (Figure 5B; Table S2; Supplementary Data 1). MVA1 and *N*-terminal derivatives are distributed widely across the chemical space (Figure 5C) with MVA1 derivatives forming at least three distinct clusters.

Inspection of the 76 derivatives that did not complete the initial parameterization and AMBER-based filtering workflow did not identify a single structural feature that consistently accounted for the failures. The unsuccessful cases were chemically diverse, and failures could not be attributed to a specific functional group or structural class based on this analysis.

Based on the single-frame MM/GBSA ranking, the top 50 derivatives from each campaign were continued from the previously minimized structures through heating and equilibration, followed by 5 ns MD. All 100 selected systems completed the 5 ns MD without failures. MM/GBSA calculations were performed over the final 2 ns of each trajectory (Supplementary Data 1). These calculations were not intended to compare or prioritize the binding properties of the derivatives in the present study, but to demonstrate that systems containing residues parameterized through TopFor could proceed through MD and trajectory-based energetic analysis in AMBER.

### Application of TopFor in a Structure-Based Peptide Discovery Workflow

To demonstrate the integration of TopFor into a structure-based peptide discovery workflow, the *E. coli* DnaN–griselimycin (GM) complex (PDB ID: 8CIX^15^) was further investigated to generate candidate peptides. A 1 µs MD simulation using TopFor-generated parameters for the noncanonical amino acids of GM was performed, and superposition of the representative MD structure with the experimental crystal structure after alignment on the DnaN backbone gave backbone RMSDs of 1.69 Å for DnaN and 1.87 Å for GM (Figure S8). Per-residue MM/GBSA decomposition performed over the final 200 ns (800–1000 ns) revealed differential contributions of individual GM residues to DnaN binding, with LEU5 showing the strongest favorable contribution (Figures S9 and S10). Based on the structural and energetic analyses, the N-terminal ACE1–LEU5 region was retained as the scaffold for peptide design.

A focused library of 89 GM-derived candidates was generated (Supplementary Data 1), comprising 61 tetrapeptides, 21 pentapeptides, and 7 *N*-terminally fatty-acid-modified derivatives with different chain lengths. The candidates were docked into the *E. coli* DnaN binding site and subjected to 10 ns MD simulations. MM/GBSA calculations were used during successive stages of computational refinement, and candidates were prioritized within their respective derivative classes based on the calculated binding energies together with structural inspection of their predicted binding poses and interactions with DnaN. Six candidates (EFB1–EFB6) were selected for synthesis and experimental evaluation.

Exploratory SPR measurements were performed using immobilized *M. smegmatis* DnaN. Although the EFB peptides were designed using the *E. coli* DnaN–GM complex, crystal structures are available for GM bound to DnaN from *E. coli*, *M. smegmatis*, and *Mycobacterium tuberculosis (M. tuberculosis)* (PDB IDs: 8CIX^15^, 5AH2^16^, and 5AGU^16^, respectively), demonstrating that the parent GM scaffold can engage DnaN from each of these species. *M. smegmatis* DnaN was therefore used as a mycobacterial DnaN system for exploratory SPR evaluation. In the initial exploratory measurement, GM at 100 nM produced a positive SPR response (54.0 RU; Table S6). Initial screening at 500 µM showed positive responses for EFB2 and EFB4–EFB6, whereas EFB1 and EFB3 showed no clear response under the screening conditions (Table S6). EFB4–EFB6 were evaluated from 62.5 µM to 1 mM and showed concentration-dependent increases in SPR response (Figure 6). Because the measurements did not permit reliable determination of equilibrium dissociation constants, the SPR results were interpreted qualitatively rather than as quantitative affinity measurements. No detectable antibacterial activity was observed for the EFB2-6 peptides against *Staphylococcus aureus* Newman and *E. coli* ΔacrB (JW0451-2) at concentrations up to 64 µM or against *M. tuberculosis* H37Ra at concentrations up to 128 µM (Table S5).

**Figure 6.**
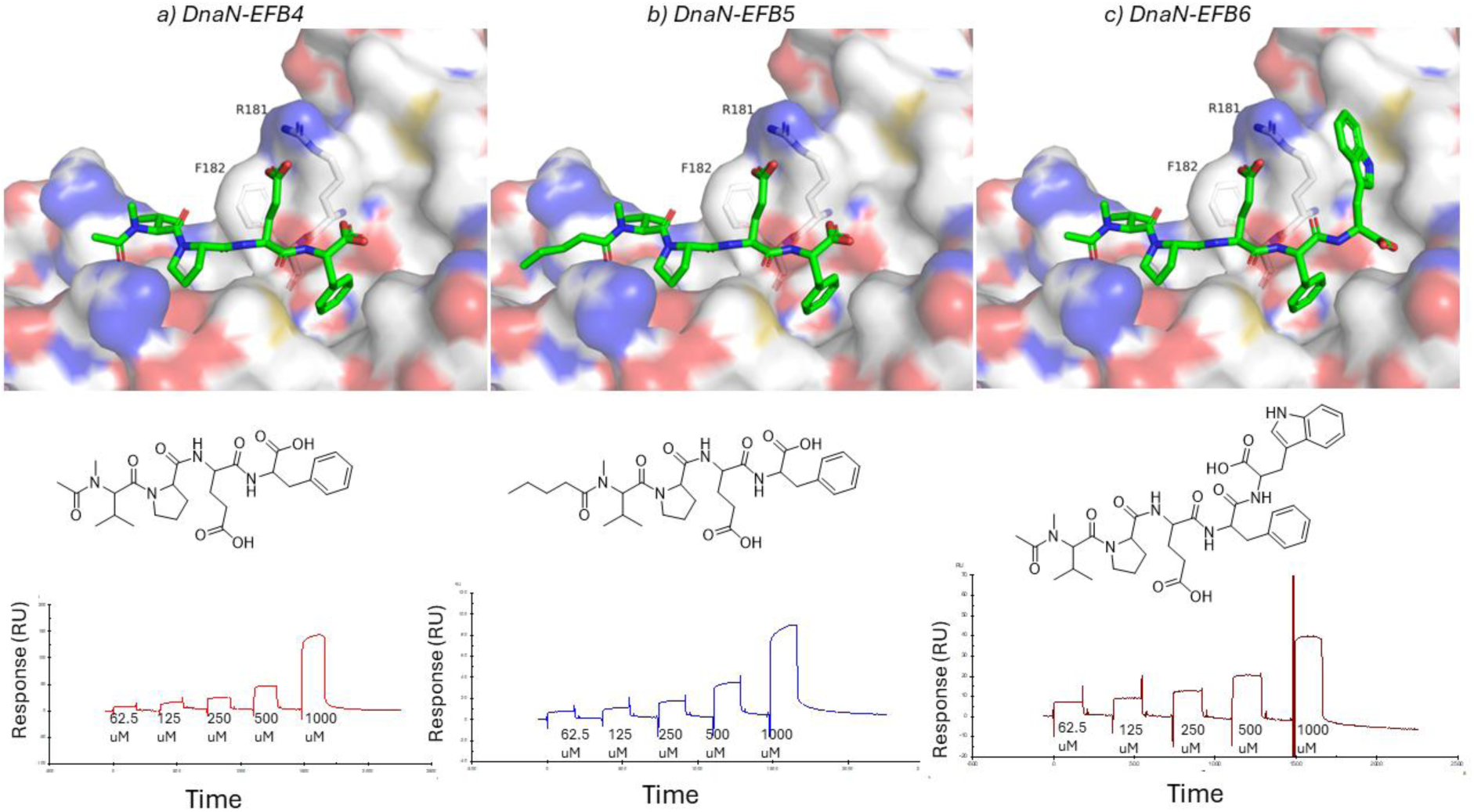
Docking poses and SPR evaluation of EFB4–EFB6 against *M. smegmatis* DnaN. (a–c) Docking poses, chemical structures, and SPR sensorgrams of EFB4 (a), EFB5 (b), and EFB6 (c). The peptides were docked into the binding site of *M. smegmatis* DnaN (PDB ID: 5AH2). DnaN is shown as an electrostatic surface, with positively charged, negatively charged, and neutral regions shown in blue, red, and white, respectively. The docked peptides are shown as sticks with carbon atoms in green, nitrogen atoms in blue, and oxygen atoms in red. The corresponding two-dimensional chemical structure of each peptide is shown below the docking pose. SPR sensorgrams obtained using immobilized *M. smegmatis* DnaN are shown at the bottom for peptide concentrations ranging from 62.5 µM to 1000 µM.

Following the SPR evaluation, EFB1–EFB6 were docked to *M. smegmatis* DnaN (PDB ID: 5AH2^16^) to examine structural features associated with the observed SPR response pattern (Figure 6 and Figure S11). The predicted binding poses showed that the residues corresponding to LEU5 of GM occupied the same pocket in all six EFB peptides. EFB2 and EFB4–EFB6 contained Phe at this position and produced positive SPR responses, whereas EFB1 and EFB3 contained Gln and Leu, respectively, and showed no clear response. At the adjacent variable position, EFB4–EFB6 contained Glu oriented toward Arg181 in the predicted complexes, whereas EFB2 contained Met. In EFB1, Arg occupied this region near Arg181, suggesting a potentially unfavorable local electrostatic environment, while EFB3 contained Tyr. These predicted interactions provide a possible structural explanation for the observed SPR response pattern but require further experimental validation.

This application demonstrates the utility of TopFor as the parameter-generation step within a workflow extending to MD-guided scaffold selection, computational peptide design, and experimental evaluation. Of the six computationally prioritized peptides, EFB2 and EFB4–EFB6 produced positive responses in the initial SPR screen, with EFB4–EFB6 showing concentration-dependent responses.

## Discussion

TopFor was developed to reduce the manual preparation required for MD simulations of peptides containing ncAAs in AMBER. Here, the workflow generated AMBER-compatible residue definitions across different peptide architectures and was applied repeatedly to large libraries of chemically modified peptides.

The MDM2–p53 benchmark provided a controlled comparison between chemically identical residues represented by native AMBER definitions or parameters prepared through TopFor. AM1-BCC, ABCG2, and RESP produced comparable structural and conformational behavior to the native AMBER representation, while the lower computational cost of AM1-BCC and ABCG2 supported their use in applications involving multiple ncAAs. The tested scope was extended to the head-to-tail cyclic cyclosporin A peptide and the cyclic depsipeptide CGM. In CGM, residues directly involved in the sidechain-to-backbone ester linkage required manual parameterization, defining a current limitation of the automated workflow.

Existing approaches cover different aspects of ncAA parameterization and differ in force-field ecosystem, molecular scope, computational requirements, and degree of automation (Table S4). TopFor does not introduce a new force field or parameterization model, but integrates established AMBER-based procedures into a peptide-oriented workflow supporting single-residue, batch, and whole-peptide inputs together with multiple charge-derivation approaches. Whole-peptide processing enables ncAAs and their positions within a peptide chain to be identified directly from molecular connectivity, whereas batch processing enables the same procedure to be applied to large numbers of chemically distinct residues. The validation presented here evaluates the automated implementation of established parameterization procedures and the use of the resulting residue definitions in AMBER simulations; it does not independently assess the accuracy of the underlying force fields or charge models for individual ncAAs.

At library scale, 1,138 of 1,214 submitted CGM derivatives (93.7%) successfully completed the end-to-end initial workflow comprising TopFor parameterization and subsequent AMBER-based filtering. All 100 derivatives selected for simulation completed 5 ns production MD and trajectory-based MM/GBSA analysis. These calculations were intended to assess whether systems parameterized through TopFor could progress through a high-throughput downstream AMBER workflow, rather than to identify optimized CGM derivatives. The current implementation primarily supports α-amino-acid residues with conventional peptide-bond connectivity. Future extensions could include support for ester, thioether, and other non-amide linkages, as well as β- and γ-amino-acid backbones.

The DnaN–GM application extended the evaluation of TopFor from parameter generation for predefined systems to its use as the parameter-generation step within a structure-based peptide design workflow. Four of six computationally prioritized peptides produced positive responses in the initial SPR screen, with EFB4–EFB6 showing concentration-dependent responses, although the measurements did not permit quantitative affinity determination. None of the tested peptides showed detectable whole-cell antibacterial activity within the investigated concentration ranges. This application demonstrated that residue parameters generated by TopFor enabled AMBER MD simulations and MM/GBSA analyses within a computational peptide-design workflow that led to experimental evaluation of selected candidates.

In summary, TopFor provides an automated and scalable route for preparing AMBER-compatible residue definitions for ncAA-containing peptides. Its main contribution is the integration of single-residue, batch, and whole-peptide processing with multiple charge-derivation options in a workflow designed for repeated preparation of chemically modified peptides. Across the systems examined here, TopFor-generated residue parameters enabled ncAA-containing peptides to be incorporated into downstream AMBER simulations and analyses, including library-scale computational screening and structure-based peptide design workflows.

## Data and Code Availability

Compound-level data generated in the high-throughput CGM derivatization campaigns and GM-derived peptide-design workflow are provided in Supplementary Data 1. TopFor is released as open-source software, together with installation instructions, the full set of command-line options, and example workflows, at https://github.com/kalininalab/TopFor.

## Supporting Information

Supplementary methods, parameterization and validation results, additional molecular dynamics analyses, docking results, SPR screening data, supplementary tables, and Figures S1–S11 (PDF).

Compound-level data from the high-throughput CGM derivatization campaigns and GM-derived peptide-design workflow, including UMAP coordinates, post-docking minimization and single-frame MM/GBSA results, trajectory-based MM/GBSA results for derivatives selected for 5 ns molecular dynamics simulations, and data underlying the GM-derived peptide-design and SPR evaluation workflows (Supplementary Data 1, XLSX).

## Author Contributions

E.F.B., O.V.K and K.A.J.B. conceived the study. E.F.B. designed the computational workflow, performed the computational analyses, and supervised the development and validation of TopFor. H.G. developed and implemented TopFor and contributed to its validation. C.B. supervised the software development and contributed to methodological development. E.F.B. and S.R. performed the SPR experiments, which were supervised by R.M. O.V.K. and K.A.J.B. supervised the study. E.F.B. and H.G. wrote the original draft. E.F.B., C.B., O.V.K. and K.A.J.B. revised the manuscript with input from all authors.

## Acknowledgment

The authors thank Viktoria George for performing the MIC assays, and Andreas Keller for scientific advice.

