## Supplementary Information for "TopFor: Automated Preparation of AMBER-Compatible Parameters for Noncanonical Amino Acids in Peptides"

**Affiliations:**

#### Supporting Information

**Supplementary Table**

**Table S1. RMSDs of representative MDM2–p53 structures relative to the experimental crystal structure (PDB ID: 1YCR).**

| Parameterization | Replicate | MDM2 backbone RMSD (Å) | p53 backbone RMSD (Å) |
|---|---|---|---|
| STANDARD | R1 | 1.234 | 1.529 |
| STANDARD | R2 | 1.163 | 1.522 |
| STANDARD | R3 | 1.260 | 1.329 |
| AM1-BCC | R1 | 1.796 | 1.535 |
| AM1-BCC | R2 | 1.169 | 1.272 |
| AM1-BCC | R3 | 1.593 | 1.231 |
| ABCG2 | R1 | 1.310 | 1.222 |
| ABCG2 | R2 | 1.359 | 1.264 |
| ABCG2 | R3 | 1.310 | 1.482 |
| RESP | R1 | 1.060 | 1.623 |
| RESP | R2 | 1.152 | 1.413 |
| RESP | R3 | 1.264 | 1.732 |

Representative MD structures were aligned to the experimental crystal structure using the MDM2 backbone. p53 backbone RMSDs were subsequently calculated for residues 86–98 using the corresponding N, Cα, and C atoms without additional fitting of the peptide.

**Table S2. Summary of the high-throughput cyclohexylgriselimycin (CGM) derivatization workflows used to evaluate the scalability of TopFor.**

| Workflow step | MVA1 substitution campaign | N-terminal derivatization campaign |
|---|---|---|
| Initial building-block library | 3,194 α-amino acids | 75,132 carboxylic acids |
| After physicochemical filtering | 1,620 | 4,486 |
| Docking protocol | Core-constrained Glide | Core-constrained Glide |
| Docking score cutoff | ≤ −6 kcal $mol^{-1}$ | ≤ −7 kcal $mol^{-1}$ |
| Peptide derivatives selected after docking | 521 | 693 |
| Completed TopFor parameterization and initial AMBER filtering | 498 | 640 |
| Not complete | 23 | 53 |
| Initial workflow completion rate (%) | 95.6% | 92.4% |
| Post-docking refinement (AMBER) | 498 | 640 |
| MM/GBSA rescoring | 498 | 640 |

| Workflow step | MVA1 substitution campaign | N-terminal derivatization campaign |
|---|---|---|
| Candidates selected for 5 ns MD | 50 | 50 |
| MM/GBSA rescoring | 50 | 50 |

**Table S3. Summary of the validation and application systems used to evaluate the TopFor workflow.** The selected systems represent increasing levels of peptide structural and chemical complexity and were used for distinct purposes, including controlled comparison with native AMBER residue definitions, application to a head-to-tail cyclic peptide containing multiple ncAAs, application to a cyclic depsipeptide containing a nonstandard ester linkage, and evaluation of library-scale parameterization and subsequent molecular simulation in AMBER. The parameterization mode, charge-derivation method, simulation protocol, and primary purpose of each application are summarized.

| System | Architecture | ncAAs processed | TopFor mode | Charge | Simulation | Purpose |
|---|---|---|---|---|---|---|
| MDM2–p53 | Linear | Canonical residues treated as ncAAs | Whole peptide | AM1-BCC / ABCG2 / RESP | 3 × 100 ns each | Controlled benchmark |
| Cyclophilin–cyclosporin A | Head-to-tail cyclic | 9 positions / 6 types | Batch | ABCG2 | 1 × 100 ns | Multi-ncAA cyclic application |
| DnaN–CGM | Cyclic depsipeptide | 4 ncAAs by TopFor + NZC/Gly ester-linked residues manually | Single | AM1-BCC | 1 × 100 ns | Complex chemistry/application boundary |
| CGM libraries | Derivative libraries | 1,214 submitted; 1,138 completed initial workflow; 100 selected for 5-ns MD | Batch | AM1-BCC | 1 × 5 ns for 100 compounds | High-throughput application |
| DnaN–GM / EFB peptides | Linear GM-derived peptides | GM ncAAs; MVA in 89 EFB derivatives | Single | AM1-BCC | 1 × 1000 ns (GM); 1 × 10 ns (89 designed derivatives) | Structure-based peptide discovery application |

**Table S4. Comparison of TopFor with existing approaches for molecular parameterization and noncanonical amino acid preparation.** The approaches are compared according to their primary parameterization target, force-field and molecular simulation ecosystem, software requirements, charge-derivation strategy, requirement for quantum-mechanical calculations, supported input modes, and demonstrated applicability to library-scale parameterization of ncAA-containing peptides.

| Feature | ACPYPE | CGenFF / CHARMM36 ncAA extension | NCAP | drFrankenstein | Wijesiri–Brown | Pep2MARS | TopFor |
|---|---|---|---|---|---|---|---|
| Primary parameterization target | Small molecules / ligands | Small molecules; extended to 333 ncAAs | Individual ncAAs for peptides/proteins | Individual ncAAs for peptides/proteins | Individual ncAAs for cyclic peptide modeling | ncAAs and covalently connected residue blocks in cyclic peptides/protein–peptide system | ncAAs in peptides |

| Feature | ACPYPE | CGenFF / CHARMM36 ncAA extension | NCAP | drFrankenstein | Wijesiri–Brown | Pep2MARS | TopFor |
|---|---|---|---|---|---|---|---|
| Force-field / MD ecosystem | GAFF/GAFF2-derived; outputs for AMBER, GROMACS, CHARMM, CNS | CHARMM36 / CGenFF | CHARMM | AMBER | AMBER + Rosetta | AMBER / GAFF; AMBER topology can be converted for GROMACS | AMBER |
| Main software / tools | ACPYPE + Antechamber | CGenFF / ParamChem + QM calculations | NCAP + NWChem + ChargeMol | drFrankenstein + QM/AmberTools workflow | Custom scripts + GAMESS + PyRESP + AmberTools + Rosetta | Pep2MARS + AmberTools + Psi4; web-based implementation | TopFor + AmberTools; xTB/ORCA/Multiwfn for RESP |
| Charge derivation | AM1-BCC via Antechamber* | CGenFF charges assigned by analogy; ncAA FF development uses QM-optimized charges | DDEC6 recommended; ChargeMol step optional | RESP / RESP2 | Ensemble RESP | RESP; AM1-BCC discussed as an alternative | AM1-BCC, ABCG2, Gasteiger, RESP |
| QM calculations required for ncAA parameterization | No for standard AM1-BCC workflow | Yes for development/refinement of new ncAA parameters | Yes | Yes | Yes | Yes for the reported RESP workflow | No for AM1-BCC/ABCG2/Gasteiger; Yes for RESP |
| Single ncAA parameterization | Not designed specifically for ncAAs | Yes, during ncAA force-field development | Yes | Yes | Yes | Yes, as monoresidue blocks within peptide systems | Yes |
| Dedicated batch ncAA parameterization | No | No dedicated peptide-library mode reported | No dedicated library mode reported | No dedicated library mode reported | No dedicated library mode reported | No dedicated residue-library mode reported | Yes |
| Whole-peptide input with automatic ncAA processing | No | No | No | No | No | Yes | Yes |
| Library-scale ncAA peptide application demonstrated | No; large-scale application demonstrated for small molecules | 333 ncAA parameter set developed, but not peptide-library parameterization/screening | No | No | No | No comparable demonstration reported | Yes; >1,000 CGM derivatives |

**Table S5. Antibacterial activity of EFB2–EFB6.** Minimum inhibitory concentrations (MICs) were determined against *Staphylococcus aureus Newman*, *Escherichia coli ΔacrB (JW0451-2),* and *Mycobacterium tuberculosis* H37Ra. Values are reported in μM from two replicate measurements. Ciprofloxacin, vancomycin, and rifampicin were included as antibacterial reference compounds. n.d., not determined.

| Compound | *S. aureus Newman [µM]* | *E. coli ΔacrB (JW0451-2) [µM]* | *M. tuberculosis H37Ra [µM]* |
|---|---|---|---|
| **EFB2** | > 64 | > 64 | > 128 |
| **EFB3** | > 64 | > 64 | > 128 |
| **EFB4** | > 64 | > 64 | > 128 |
| **EFB5** | > 64 | > 64 | > 128 |
| **EFB6** | > 64 | > 64 | > 128 |
| Ciprofloxacin | | 0.01 | |
| Vancomycin | 2 | | |
| Rifampicin | | | 0.06 |

**Table S6. Exploratory SPR screening of EFB1–EFB6 at 500 µM and griselimycin (GM) at 100 nM against Mycobacterium smegmatis DnaN.** SPR responses are reported as response units (RU) after reference subtraction (Fc = 2–1). The measurements were performed as a single exploratory screen to assess detectable binding responses and were not used for quantitative determination of equilibrium dissociation constants.

| Cycle | Curve | Concentration | Response (RU) |
|---|---|---|---|
| **1** | **Fc=2-1** | **EFB1_500uM** | **-4.6** |
| **1** | **Fc=2-1** | **EFB2_500uM** | **11.3** |
| **1** | **Fc=2-1** | **EFB3_500uM** | **1.8** |
| **1** | **Fc=2-1** | **EFB4_500uM** | **47.5** |
| **1** | **Fc=2-1** | **EFB5_500uM** | **48.4** |
| **1** | **Fc=2-1** | **EFB6_500uM** | **33.1** |
| **1** | **Fc=2-1** | **GM_100nM** | **54.0** |

**Table S7. Sequences and synthesis details of EFB1–EFB6.**

| Peptide | Sequence | *N*-terminal modification | Purity | Quantity |
|---|---|---|---|---|
| EFB1 | Ac-{NMe-Val}PRQ | Acetylation | ≥90% | 14 mg |
| EFB2 | Ac-{NMe-Val}PMF | Acetylation | ≥90% | 14 mg |
| EFB3 | Ac-{NMe-Val}PYL | Acetylation | ≥90% | 14 mg |
| EFB4 | Ac-{NMe-Val}PEF | Acetylation | ≥90% | 14 mg |
| EFB5 | Pentanoyl-{NMe-Val}PEF | Pentanoic acid | ≥90% | 14 mg |
| EFB6 | Ac-{NMe-Val}PEFW | Acetylation | ≥90% | 14 mg |

**Representative TopFor output files for N-methylvaline**

Representative intermediate and final files generated during the parameterization of N-methylvaline (MVA) are presented in Tables S8–S15. The example includes the initial residue structure, the capped structure generated for parameterization, and the AC, MC, PREPIN, FRCMOD, and LIB files generated during the TopFor workflow.

**Table S8. Input MOL2 structure of N-methylvaline (MVA) used for TopFor parameterization.**

```
@<TRIPOS>MOLECULE
8CIY.B
19 18 1 0 0
BIOPOLYMER
NO_CHARGES
@<TRIPOS>ATOM
 1 N     -15.7150  -13.5540    1.7340 N.3   1 MVA2   0.0000 BACKBONE
 2 CN    -15.9010  -13.9890    0.3360 C.3   1 MVA2   0.0000
 3 CA    -16.8170  -13.9400    2.6610 C.3   1 MVA2   0.0000 BACKBONE
 4 CB    -18.1240  -13.1710    2.4370 C.3   1 MVA2   0.0000
 5 CG2   -19.0510  -13.4180    3.6180 C.3   1 MVA2   0.0000
 6 CG1   -17.8220  -11.6950    2.2760 C.3   1 MVA2   0.0000
 7 C     -16.9750  -15.4560    2.5770 C.2   1 MVA2   0.0000 BACKBONE
 8 O     -17.9530  -15.9480    2.0330 O.2   1 MVA2   0.0000 BACKBONE
 9 HA    -16.5620  -13.6930    3.5840 H     1 MVA2   0.0000
10 HB    -18.5670  -13.4890    1.6110 H     1 MVA2   0.0000
11 HG11  -18.6540  -11.2040    2.1670 H     1 MVA2   0.0000
12 HG12  -17.3560  -11.3710    3.0650 H     1 MVA2   0.0000
13 HG13  -17.2630  -11.5620    1.4920 H     1 MVA2   0.0000
14 HG21  -19.2370  -14.3690    3.6920 H     1 MVA2   0.0000
15 HG22  -19.8830  -12.9350    3.4840 H     1 MVA2   0.0000
16 HG23  -18.6250  -13.1080    4.4350 H     1 MVA2   0.0000
17 HN1   -16.7570  -14.4410    0.2440 H     1 MVA2   0.0000
18 HN2   -15.8820  -13.2150   -0.2540 H     1 MVA2   0.0000
19 HN3   -15.1870  -14.6000    0.0850 H     1 MVA2   0.0000
@<TRIPOS>BOND
 1  1  2 1
 2  1  3 1  BACKBONE
 3  2 17 1
```

4 2 18 1

5 2 19 1

6 3 4 1

7 3 7 1 BACKBONE

8 3 9 1

9 4 5 1

10 4 6 1

11 4 10 1

12 5 14 1

13 5 15 1

14 5 16 1

15 6 11 1

16 6 12 1

17 6 13 1

18 7 8 2 BACKBONE

@<TRIPOS>SUBSTRUCTURE

1 MVA2 3 RESIDUE 4 B MVA 0

### MOE 2024.06 (io_trps.svl 2023.06)

**Table S9. Capped MOL2 structure of MVA generated during residue preparation.**

### created with PyMOL 3.1.0

@<TRIPOS>MOLECULE

8CIY.B

31 30 3

SMALL

USER_CHARGES

@<TRIPOS>ATOM

1 AC1 -14.830-12.6652.038 C.2 1 ACE1 0.597

2 AC2 -14.885-12.1023.139 O.2 1 ACE1 -0.568

3 AC3 -13.741-12.3281.043 C.3 1 ACE1 -0.366

4 AC4 -12.771-12.5341.471 H 1 ACE1 0.112

5 AC5 -13.861-12.9200.148 H 1 ACE1 0.112

6 AC6 -13.793-11.2820.779 H 1 ACE1 0.112

| | | | | | | |
|---|---|---|---|---|---|---|
| 7 | N | -15.715-13.5541.734 | N.pl3 | 2 | MVA2 | 0.000 |
| 8 | CA | -16.817-13.9402.661 | C.3 | 2 | MVA2 | 0.000 |
| 9 | C | -16.975-15.4562.577 | C.2 | 2 | MVA2 | 0.000 |
| 10 | O | -16.979-16.0191.492 | O.2 | 2 | MVA2 | 0.000 |
| 11 | CB | -18.124-13.1712.437 | C.3 | 2 | MVA2 | 0.000 |
| 12 | CG1 | -17.822-11.6952.276 | C.3 | 2 | MVA2 | 0.000 |
| 13 | CG2 | -19.051-13.4183.618 | C.3 | 2 | MVA2 | 0.000 |
| 14 | CN | -15.714-14.2670.442 | C.3 | 2 | MVA2 | 0.000 |
| 15 | H01 | -16.549-13.6533.678 | H | 2 | MVA2 | 0.000 |
| 16 | H02 | -18.617-13.5161.528 | H | 2 | MVA2 | 0.000 |
| 17 | H03 | -16.863-11.5741.773 | H | 2 | MVA2 | 0.000 |
| 18 | H04 | -18.606-11.2261.681 | H | 2 | MVA2 | 0.000 |
| 19 | H05 | -17.779-11.2233.258 | H | 2 | MVA2 | 0.000 |
| 20 | H06 | -20.015-13.7733.254 | H | 2 | MVA2 | 0.000 |
| 21 | H07 | -18.612-14.1694.274 | H | 2 | MVA2 | 0.000 |
| 22 | H08 | -19.192-12.4894.171 | H | 2 | MVA2 | 0.000 |
| 23 | H09 | -15.194-13.667-0.304 | H | 2 | MVA2 | 0.000 |
| 24 | H10 | -15.207-15.2250.555 | H | 2 | MVA2 | 0.000 |
| 25 | H11 | -16.742-14.4360.119 | H | 2 | MVA2 | 0.000 |
| 26 | NM1 | -17.105-16.1473.658 | N.pl3 | 3 | NME3 | -0.416 |
| 27 | NM2 | -17.255-17.5903.584 | C.3 | 3 | NME3 | -0.149 |
| 28 | NM3 | -17.102-15.6854.557 | H | 3 | NME3 | 0.272 |
| 29 | NM4 | -18.286-17.8443.388 | H | 3 | NME3 | 0.098 |
| 30 | NM5 | -16.640-17.9862.789 | H | 3 | NME3 | 0.098 |
| 31 | NM6 | -16.955-18.0424.518 | H | 3 | NME3 | 0.098 |

@<TRIPOS>BOND

1 1 2 2

2 1 3 1

3 3 4 1

4 3 5 1

5 3 6 1

6 1 7 1

7 7 8 1

8 7 14 1

9 8 9 1

10 8 11 1

11 8 15 1

12 9 10 2

13 9 26 1

14 11 12 1

15 11 13 1

16 11 16 1

17 12 17 1

18 12 18 1

19 12 19 1

20 13 20 1

21 13 21 1

22 13 22 1

23 14 23 1

24 14 24 1

25 14 25 1

26 26 27 1

27 26 28 1

28 27 29 1

29 27 30 1

30 27 31 1

@<TRIPOS>SUBSTRUCTURE

| | | | | |
|---|---|---|---|---|
| 1 | ACE1 | 1 | GROUP1 B | ACE |
| 2 | MVA2 | 7 | GROUP1 B | MVA |
| 3 | NME3 | 26 | GROUP1 B | NME |

**Table S10. AC file of capped MVA generated following charge derivation and AMBER atom-type assignment.**

CHARGE 0.00 ( 0 )

Formula: H18 C9 N2 O2

ATOM 1 CC1 ACE 1 -14.830 -12.665 2.038 0.661035 C

ATOM 2 OC2 ACE 1 -14.885 -12.102 3.139 -0.628165 O

ATOM 3 CC3 ACE 1 -13.741 -12.328 1.043 -0.167165 CT

ATOM 4 HC4 ACE 1 -12.771 -12.534 1.471 0.068968 HC

ATOM 5 HC5 ACE 1 -13.861 -12.920 0.148 0.068968 HC

ATOM 6 HC6 ACE 1 -13.793 -11.282 0.779 0.068968 HC

ATOM 7 N MVA 2 -15.715 -13.554 1.734 -0.468865 N

ATOM 8 CA MVA 2 -16.817 -13.940 2.661 0.038635 CX

ATOM 9 C MVA 2 -16.975 -15.456 2.577 0.635035 C

ATOM 10 O MVA 2 -16.979 -16.019 1.492 -0.620165 O

ATOM 11 CB MVA 2 -18.124 -13.171 2.437 -0.064765 CT

ATOM 12 CG1 MVA 2 -17.822 -11.695 2.276 -0.093165 CT

ATOM 13 CG2 MVA 2 -19.051 -13.418 3.618 -0.093165 CT

ATOM 14 CN MVA 2 -15.714 -14.267 0.442 0.077235 CT

ATOM 15 H01 MVA 2 -16.549 -13.653 3.678 0.094635 H1

ATOM 16 H02 MVA 2 -18.617 -13.516 1.528 0.073635 HC

ATOM 17 H03 MVA 2 -16.863 -11.574 1.773 0.039802 HC

ATOM 18 H04 MVA 2 -18.606 -11.226 1.681 0.039802 HC

ATOM 19 H05 MVA 2 -17.779 -11.223 3.258 0.039802 HC

ATOM 20 H06 MVA 2 -20.015 -13.773 3.254 0.039802 HC

ATOM 21 H07 MVA 2 -18.612 -14.169 4.274 0.039802 HC

ATOM 22 H08 MVA 2 -19.192 -12.489 4.171 0.039802 HC

ATOM 23 H09 MVA 2 -15.194 -13.667 -0.304 0.045635 H1

ATOM 24 H10 MVA 2 -15.207 -15.225 0.555 0.045635 H1

ATOM 25 H11 MVA 2 -16.742 -14.436 0.119 0.045635 H1

ATOM 26 NM1 NME 3 -17.105 -16.147 3.658 -0.568965 N

ATOM 27 CM2 NME 3 -17.255 -17.590 3.584 0.078235 CT

ATOM 28 HM3 NME 3 -17.102 -15.685 4.557 0.335435 H

ATOM 29 HM4 NME 3 -18.286 -17.844 3.388 0.042635 H1

ATOM 30 HM5 NME 3 -16.640 -17.986 2.789 0.042635 H1

ATOM 31 HM6 NME 3 -16.955 -18.042 4.518 0.042635 H1

```
BOND  1  1  2  2  CC1 OC2
BOND  2  1  3  1  CC1 CC3
BOND  3  3  4  1  CC3 HC4
BOND  4  3  5  1  CC3 HC5
BOND  5  3  6  1  CC3 HC6
BOND  6  1  7  1  CC1  N
BOND  7  7  8  1   N  CA
BOND  8  7 14  1   N  CN
BOND  9  8  9  1  CA   C
BOND 10  8 11  1  CA  CB
BOND 11  8 15  1  CA H01
BOND 12  9 10  2   C   O
BOND 13  9 26  1   C NM1
BOND 14 11 12  1  CB CG1
BOND 15 11 13  1  CB CG2
BOND 16 11 16  1  CB H02
BOND 17 12 17  1 CG1 H03
BOND 18 12 18  1 CG1 H04
BOND 19 12 19  1 CG1 H05
BOND 20 13 20  1 CG2 H06
BOND 21 13 21  1 CG2 H07
BOND 22 13 22  1 CG2 H08
BOND 23 14 23  1  CN H09
BOND 24 14 24  1  CN H10
BOND 25 14 25  1  CN H11
BOND 26 26 27  1 NM1 CM2
BOND 27 26 28  1 NM1 HM3
BOND 28 27 29  1 CM2 HM4
BOND 29 27 30  1 CM2 HM5
BOND 30 27 31  1 CM2 HM6
```

**Table S11. Main-chain definition (MC) file generated for MVA and used for PREPIN residue-template generation.**

```
HEAD_NAME N
TAIL_NAME C
MAIN_CHAIN CA
OMIT_NAME CC1
OMIT_NAME OC2
OMIT_NAME CC3
OMIT_NAME HC4
OMIT_NAME HC5
OMIT_NAME HC6
OMIT_NAME NM1
OMIT_NAME CM2
OMIT_NAME HM3
OMIT_NAME HM4
OMIT_NAME HM5
OMIT_NAME HM6
PRE_HEAD_TYPE C
POST_TAIL_TYPE N
CHARGE 0.0
```

**Table S12. PREPIN residue template generated for MVA.**

```
   0   0   2
This is a remark line
molecule.res
MVA   INT  0
CORRECT     OMIT DU   BEG
  0.0000
   1  DUMM  DU    M    0  -1  -2     0.000      .0       .0      .00000
   2  DUMM  DU    M    1   0  -1     1.449      .0       .0      .00000
   3  DUMM  DU    M    2   1   0     1.523   111.21      .0      .00000
   4  N     N     M    3   2   1     1.540   111.208  -180.000  -0.460816
   5  CN    CT    3    4   3   2     1.476    90.039    61.108  0.078561
```

```
 6 H09  H1   E   5  4  3    1.089  109.496  120.404 0.046418
 7 H10  H1   E   5  4  3    1.090  109.529 -119.564 0.046418
 8 H11  H1   E   5  4  3    1.091  109.481    0.451 0.046418
 9 CA   CX   M   4  3  2    1.491   42.340  -67.393 0.039298
10 CB   CT   3   9  4  3    1.533  114.173  -26.491 -0.063653
11 CG1  CT   3  10  9  4    1.515  109.529  -45.516 -0.091566
12 H03  HC   E  11 10  9    1.090  109.430   33.817 0.040485
13 H04  HC   E  11 10  9    1.090  109.495  153.795 0.040485
14 H05  HC   E  11 10  9    1.090  109.503  -86.149 0.040485
15 CG2  CT   3  10  9  4    1.522  108.942 -167.377 -0.091566
16 H06  HC   E  15 10  9    1.090  109.422 -126.582 0.040485
17 H07  HC   E  15 10  9    1.090  109.497   -6.624 0.040485
18 H08  HC   E  15 10  9    1.090  109.521  113.476 0.040485
19 H02  HC   E  10  9  4    1.090  110.407   73.347 0.074899
20 H01  H1   E   9  4  3    1.090  109.279 -140.403 0.096260
21 C    C    M   9  4  3    1.527  107.422  100.564 0.645936
22 O    O    E  21  9  4    1.222  120.437  -44.978 -0.609519
LOOP
IMPROPER
  -M  CN   N  CA
  CA  +M   C   O
DONE
STOP
```

**Table S13. ff19SB FRCMOD file generated for MVA.**

```
Remark line goes here
MASS
C  12.010      0.616
O  16.000      0.434
CT 12.010      0.878
HC 1.008       0.135
N  14.010      0.530
CX 12.010      0.360
```

H1 1.008 0.135

H 1.008 0.161

BOND

C -O 570.00 1.229

C -CT 317.00 1.522

C -N 490.00 1.335

CT-HC 340.00 1.090

CX-N 337.00 1.449

CT-N 337.00 1.449

C -CX 317.00 1.522

CT-CX 310.00 1.526

CX-H1 340.00 1.090

CT-CT 310.00 1.526

CT-H1 340.00 1.090

H -N 434.00 1.010

ANGLE

C -CT-HC 50.000 109.500

C -N -CX 50.000 121.900

C -N -CT 50.000 121.900

CT-C -O 80.000 120.400

N -C -O 80.000 122.900

CT-C -N 70.000 116.600

HC-CT-HC 35.000 109.500

C -CX-N 63.000 110.100

CT-CX-N 80.000 109.700

H1-CX-N 50.000 109.500

H1-CT-N 50.000 109.500

CT-N -CX 50.000 118.000

CX-C -O 80.000 120.400

CX-C -N 70.000 116.600

CT-CT-CX 40.000 109.500

CX-CT-HC 50.000 109.500

C -CX-CT 63.000 111.100

C -CX-H1 50.000 109.500

C -N -H 50.000 120.000

CT-CX-H1 50.000 109.500

CT-CT-HC 50.000 109.500

CT-CT-CT 40.000 109.500

H1-CT-H1 35.000 109.500

CT-N -H 50.000 118.040

DIHE

C -CX-N -C 1 0.000 0.000 -4.000

C -CX-N -C 1 0.420 0.000 -3.000

C -CX-N -C 1 0.270 0.000 -2.000

C -CX-N -C 1 0.000 0.000 1.000

CT-CX-N -C 1 0.000 0.000 -4.000

CT-CX-N -C 1 0.400 0.000 -3.000

CT-CX-N -C 1 2.000 0.000 -2.000

CT-CX-N -C 1 2.000 0.000 1.000

H1-CX-N -C 6 0.000 0.000 2.000

H1-CT-N -C 6 0.000 0.000 2.000

O -C -CT-HC 1 0.800 0.000 -1.000

O -C -CT-HC 1 0.000 0.000 -2.000

O -C -CT-HC 1 0.080 180.000 3.000

O -C -N -CX 4 10.000 180.000 2.000

O -C -N -CT 4 10.000 180.000 2.000

CT-C -N -CX 4 10.000 180.000 2.000

CT-C -N -CT 4 10.000 180.000 2.000

N -C -CT-HC 6 0.000 0.000 2.000

O -C -CX-N 6 0.000 0.000 2.000

N -C -CX-N 1 0.000 0.000 -4.000

N -C -CX-N 1 0.550 180.000 -3.000

N -C -CX-N 1 1.580 180.000 -2.000

N -C -CX-N   1   0.450      180.000        1.000

CT-CT-CX-N   9   1.400       0.000        3.000

HC-CT-CX-N   9   1.400       0.000        3.000

H1-CT-N -CX  6   0.000       0.000        2.000

CX-C -N -CT  4  10.000      180.000        2.000

CX-C -N -H   4  10.000      180.000        2.000

CX-CT-CT-HC  1   0.160       0.000        3.000

CT-CT-CX-C   9   1.400       0.000        3.000

HC-CT-CX-C   9   1.400       0.000        3.000

O -C -N -H   1   2.500      180.000       -2.000

O -C -N -H   1   2.000       0.000        1.000

O -C -CX-CT  6   0.000       0.000        2.000

N -C -CX-CT  1   0.000       0.000       -4.000

N -C -CX-CT  1   0.400       0.000       -3.000

N -C -CX-CT  1   0.200       0.000       -2.000

N -C -CX-CT  1   0.200       0.000        1.000

CT-CT-CT-HC  1   0.160       0.000        3.000

C -CX-N -CT  6   0.000       0.000        2.000

CT-CX-N -CT  6   0.000       0.000        2.000

H1-CX-N -CT  6   0.000       0.000        2.000

O -C -CX-H1  1   0.800       0.000       -1.000

O -C -CX-H1  1   0.000       0.000       -2.000

O -C -CX-H1  1   0.080      180.000        3.000

N -C -CX-H1  6   0.000       0.000        2.000

CT-CT-CX-H1  9   1.400       0.000        3.000

HC-CT-CX-H1  9   1.400       0.000        3.000

HC-CT-CT-HC  1   0.150       0.000        3.000

H1-CT-N -H   6   0.000       0.000        2.000

IMPROPER

CT-N -C -O       10.5       180.0       2.0       Using general improper torsional angle  X- X- C- O, penalty score=  6.0)

C -CT-N -CX       1.1       180.0       2.0       Using the default value

CX-N -C -O 10.5 180.0 2.0 Using general improper torsional angle X- X- C- O, penalty score= 6.0)

C -CT-N -H 1.1 180.0 2.0

NONBON

C 1.9080 0.0860

O 1.6612 0.2100

CT 1.9080 0.1094

HC 1.4870 0.0157

N 1.8240 0.1700

CX 1.9080 0.1094

H1 1.3870 0.0157

H 0.6000 0.0157

**Table S14. GAFF2 FRCMOD file generated for MVA.**

Remark line goes here

MASS

C 12.010 0.616 same as c

O 16.000 0.434 same as o

CT 12.010 0.878 same as c3

HC 1.008 0.135 same as hc

N 14.010 0.530 same as n

CX 12.010 0.878 same as c3

H1 1.008 0.135 same as h1

H 1.008 0.161 same as hn

BOND

C -O 590.60 1.219 same as c- o, penalty score= 0.0

C -CT 234.85 1.527 same as c-c3, penalty score= 0.0

C -N 317.46 1.397 same as c- n, penalty score= 0.0

CT-HC 345.25 1.096 same as c3-hc, penalty score= 0.0

CX-N 254.46 1.464 same as c3- n, penalty score= 0.0

CT-N 254.46 1.464 same as c3- n, penalty score= 0.0

C -CX 234.85 1.527 same as c-c3, penalty score= 0.0

CT-CX 228.89 1.535 same as c3-c3, penalty score= 0.0

CX-H1  344.22   1.097      same as c3-h1, penalty score=  0.0

CT-CT  228.89   1.535      same as c3-c3, penalty score=  0.0

CT-H1  344.22   1.097      same as c3-h1, penalty score=  0.0

H -N   527.30   1.013      same as hn- n, penalty score=  0.0

ANGLE

C -CT-HC   43.630    108.950   same as c -c3-hc, penalty score=  0.0

C -N -CX   77.240    118.830   same as c -n -c3, penalty score=  0.0

C -N -CT   77.240    118.830   same as c -n -c3, penalty score=  0.0

CT-C -O   76.450    122.900   same as c3-c -o , penalty score=  0.0

N -C -O   86.680    122.570   same as n -c -o , penalty score=  0.0

CT-C -N   65.190    117.160   same as c3-c -n , penalty score=  0.0

HC-CT-HC   35.800    107.730   same as hc-c3-hc, penalty score=  0.0

C -CX-N   65.390    111.940   same as c -c3-n , penalty score=  0.0

CT-CX-N   64.950    112.780   same as c3-c3-n , penalty score=  0.0

H1-CX-N   48.090    108.760   same as h1-c3-n , penalty score=  0.0

H1-CT-N   48.090    108.760   same as h1-c3-n , penalty score=  0.0

CT-N -CX   76.210    116.750   same as c3-n -c3, penalty score=  0.0

CX-C -O   76.450    122.900   same as c3-c -o , penalty score=  0.0

CX-C -N   65.190    117.160   same as c3-c -n , penalty score=  0.0

CT-CT-CX   59.870    112.630   same as c3-c3-c3, penalty score=  0.0

CX-CT-HC   43.270    109.680   same as c3-c3-hc, penalty score=  0.0

C -CX-CT   60.360    111.430   same as c -c3-c3, penalty score=  0.0

C -CX-H1   43.740    108.370   same as c -c3-h1, penalty score=  0.0

C -N -H   48.700    117.550   same as c -n -hn, penalty score=  0.0

CT-CX-H1   43.280    109.590   same as c3-c3-h1, penalty score=  0.0

CT-CT-HC   43.270    109.680   same as c3-c3-hc, penalty score=  0.0

CT-CT-CT   59.870    112.630   same as c3-c3-c3, penalty score=  0.0

H1-CT-H1   35.640    108.550   same as h1-c3-h1, penalty score=  0.0

CT-N -H   46.100    117.680   same as c3-n -hn, penalty score=  0.0

DIHE

C -CX-N -C   1   0.390      180.000        -2.000      same as c -c3-n -c

C -CX-N -C   1   0.640       0.000         1.000      same as c -c3-n -c , penalty score=  0.0

CT-CX-N -C 1 0.650 180.000 -4.000 same as c3-c3-n -c

CT-CX-N -C 1 0.030 180.000 -3.000 same as c3-c3-n -c

CT-CX-N -C 1 2.260 0.000 1.000 same as c3-c3-n -c , penalty score= 0.0

H1-CX-N -C 1 0.000 180.000 2.000 same as h1-c3-n -c , penalty score= 0.0

H1-CT-N -C 1 0.000 180.000 2.000 same as h1-c3-n -c , penalty score= 0.0

O -C -CT-HC 1 4.030 0.000 -1.000 same as o -c -c3-hc

O -C -CT-HC 1 3.660 180.000 2.000 same as o -c -c3-hc, penalty score= 0.0

O -C -N -CX 4 10.000 180.000 2.000 same as X -c -n -X , penalty score= 0.0

O -C -N -CT 4 10.000 180.000 2.000 same as X -c -n -X , penalty score= 0.0

CT-C -N -CX 1 1.410 180.000 -2.000 same as c3-c -n -c3

CT-C -N -CX 1 2.930 0.000 -1.000 same as c3-c -n -c3

CT-C -N -CX 1 0.500 0.000 3.000 same as c3-c -n -c3, penalty score= 0.0

CT-C -N -CT 1 1.410 180.000 -2.000 same as c3-c -n -c3

CT-C -N -CT 1 2.930 0.000 -1.000 same as c3-c -n -c3

CT-C -N -CT 1 0.500 0.000 3.000 same as c3-c -n -c3, penalty score= 0.0

N -C -CT-HC 6 0.000 180.000 2.000 same as X -c -c3-X , penalty score= 0.0

O -C -CX-N 6 0.000 180.000 2.000 same as X -c -c3-X , penalty score= 0.0

N -C -CX-N 1 0.100 180.000 -1.000 same as n -c -c3-n

N -C -CX-N 1 2.120 180.000 2.000 same as n -c -c3-n , penalty score= 0.0

CT-CT-CX-N 1 0.100 0.000 3.000 same as c3-c3-c3-n , penalty score= 0.0

HC-CT-CX-N 9 1.400 0.000 3.000 same as X -c3-c3-X , penalty score= 0.0

H1-CT-N -CX 6 0.000 0.000 2.000 same as X -c3-n -X , penalty score= 0.0

CX-C -N -CT 1 1.410 180.000 -2.000 same as c3-c -n -c3

CX-C -N -CT 1 2.930 0.000 -1.000 same as c3-c -n -c3

CX-C -N -CT 1 0.500 0.000 3.000 same as c3-c -n -c3, penalty score= 0.0

CX-C -N -H 4 10.000 180.000 2.000 same as X -c -n -X , penalty score= 0.0

CX-CT-CT-HC 1 0.130 0.000 3.000 same as c3-c3-c3-hc, penalty score= 0.0

CT-CT-CX-C 1 0.000 0.000 -3.000 same as c -c3-c3-c3

CT-CT-CX-C 1 0.040 0.000 -2.000 same as c -c3-c3-c3

CT-CT-CX-C 1 0.460 180.000 1.000 same as c -c3-c3-c3, penalty score= 0.0

HC-CT-CX-C 9 1.400 0.000 3.000 same as X -c3-c3-X , penalty score= 0.0

O -C -N -H 1 2.500 180.000 -2.000 same as o -c -n -hn

O -C -N -H 1 2.000 0.000 1.000 same as o -c -n -hn, penalty score= 0.0

O -C -CX-CT 1 0.200 180.000 -2.000 same as o -c -c3-c3

O -C -CX-CT 1 0.440 180.000 -3.000 same as o -c -c3-c3

O -C -CX-CT 1 0.680 0.000 1.000 same as o -c -c3-c3, penalty score= 0.0

N -C -CX-CT 1 0.000 180.000 -4.000 same as n -c -c3-c3

N -C -CX-CT 1 0.710 180.000 2.000 same as n -c -c3-c3, penalty score= 0.0

CT-CT-CT-HC 1 0.130 0.000 3.000 same as c3-c3-c3-hc, penalty score= 0.0

C -CX-N -CT 6 0.000 0.000 2.000 same as X -c3-n -X , penalty score= 0.0

CT-CX-N -CT 6 0.000 0.000 2.000 same as X -c3-n -X , penalty score= 0.0

H1-CX-N -CT 6 0.000 0.000 2.000 same as X -c3-n -X , penalty score= 0.0

O -C -CX-H1 1 0.800 0.000 -1.000 same as o -c -c3-h1

O -C -CX-H1 1 0.080 180.000 3.000 same as o -c -c3-h1, penalty score= 0.0

N -C -CX-H1 6 0.000 180.000 2.000 same as X -c -c3-X , penalty score= 0.0

CT-CT-CX-H1 9 1.400 0.000 3.000 same as X -c3-c3-X , penalty score= 0.0

HC-CT-CX-H1 9 1.400 0.000 3.000 same as X -c3-c3-X , penalty score= 0.0

HC-CT-CT-HC 1 0.120 0.000 3.000 same as hc-c3-c3-hc, penalty score= 0.0

H1-CT-N -H 6 0.000 0.000 2.000 same as X -c3-n -X , penalty score= 0.0

IMPROPER

CT-N -C -O 1.1 180.0 2.0 Using the default value

C -CT-N -CX 1.1 180.0 2.0 Using the default value

CX-N -C -O 1.1 180.0 2.0 Using the default value

C -CT-N -H 1.1 180.0 2.0 Same as X -X -n -hn, penalty score= 6.0 (use general term))

NONBON

C 1.8606 0.0988 same as c

O 1.7107 0.1463 same as o

CT 1.9069 0.1078 same as c3

HC 1.4593 0.0208 same as hc

N 1.7852 0.1636 same as n

CX 1.9069 0.1078 same as c3

H1 1.3593 0.0208 same as h1

H 0.6210 0.0100 same as hn

**Table S15. AMBER residue library (LIB) file generated for MVA.**

!!index array str

"MVA"

!entry.MVA.unit.atoms table str name str type int typex int resx int flags int seq int elmnt dbl chg

"CC1" "C" 0 1 131072 1 6 0.661035

"OC2" "O" 0 1 131072 2 8 -0.628165

"CC3" "CT" 0 1 131072 3 6 -0.167165

"HC4" "HC" 0 1 131072 4 1 0.068968

"HC5" "HC" 0 1 131072 5 1 0.068968

"HC6" "HC" 0 1 131072 6 1 0.068968

"N" "N" 0 1 131072 7 7 -0.468865

"CA" "CX" 0 1 131072 8 6 0.038635

"C" "C" 0 1 131072 9 6 0.635035

"O" "O" 0 1 131072 10 8 -0.620165

"CB" "CT" 0 1 131072 11 6 -0.064765

"CG1" "CT" 0 1 131072 12 6 -0.093165

"CG2" "CT" 0 1 131072 13 6 -0.093165

"CN" "CT" 0 1 131072 14 6 0.077235

"H01" "H1" 0 1 131072 15 1 0.094635

"H02" "HC" 0 1 131072 16 1 0.073635

"H03" "HC" 0 1 131072 17 1 0.039802

"H04" "HC" 0 1 131072 18 1 0.039802

"H05" "HC" 0 1 131072 19 1 0.039802

"H06" "HC" 0 1 131072 20 1 0.039802

"H07" "HC" 0 1 131072 21 1 0.039802

"H08" "HC" 0 1 131072 22 1 0.039802

"H09" "H1" 0 1 131072 23 1 0.045635

"H10" "H1" 0 1 131072 24 1 0.045635

"H11" "H1" 0 1 131072 25 1 0.045635

"NM1" "N" 0 1 131072 26 7 -0.568965

"CM2" "CT" 0 1 131072 27 6 0.078235

"HM3" "H" 0 1 131072 28 1 0.335435

"HM4" "H1" 0 1 131072 29 1 0.042635

"HM5" "H1" 0 1 131072 30 1 0.042635

"HM6" "H1" 0 1 131072 31 1 0.042635

!entry.MVA.unit.atomspertinfo table  str pname  str ptype  int ptypex  int pelmnt  dbl pchg

"CC1" "C" 0 -1 0.0

"OC2" "O" 0 -1 0.0

"CC3" "CT" 0 -1 0.0

"HC4" "HC" 0 -1 0.0

"HC5" "HC" 0 -1 0.0

"HC6" "HC" 0 -1 0.0

"N" "N" 0 -1 0.0

"CA" "CX" 0 -1 0.0

"C" "C" 0 -1 0.0

"O" "O" 0 -1 0.0

"CB" "CT" 0 -1 0.0

"CG1" "CT" 0 -1 0.0

"CG2" "CT" 0 -1 0.0

"CN" "CT" 0 -1 0.0

"H01" "H1" 0 -1 0.0

"H02" "HC" 0 -1 0.0

"H03" "HC" 0 -1 0.0

"H04" "HC" 0 -1 0.0

"H05" "HC" 0 -1 0.0

"H06" "HC" 0 -1 0.0

"H07" "HC" 0 -1 0.0

"H08" "HC" 0 -1 0.0

"H09" "H1" 0 -1 0.0

"H10" "H1" 0 -1 0.0

"H11" "H1" 0 -1 0.0

"NM1" "N" 0 -1 0.0

"CM2" "CT" 0 -1 0.0

"HM3" "H" 0 -1 0.0

"HM4" "H1" 0 -1 0.0

"HM5" "H1" 0 -1 0.0

"HM6" "H1" 0 -1 0.0

!entry.MVA.unit.boundbox array dbl

-1.000000

0.0

0.0

0.0

0.0

!entry.MVA.unit.childsequence single int

2

!entry.MVA.unit.connect array int

0

0

!entry.MVA.unit.connectivity table  int atom1x  int atom2x  int flags

1 2 2

1 3 1

1 7 1

3 4 1

3 5 1

3 6 1

7 8 1

7 14 1

8 9 1

8 11 1

8 15 1

9 10 2

9 26 1

11 12 1

11 13 1

11 16 1

12 17 1

12 18 1

12 19 1

13 20 1

13 21 1

13 22 1

14 23 1

14 24 1

14 25 1

26 27 1

26 28 1

27 29 1

27 30 1

27 31 1

!entry.MVA.unit.hierarchy table  str abovetype  int abovex  str belowtype  int belowx

"U" 0 "R" 1

"R" 1 "A" 1

"R" 1 "A" 2

"R" 1 "A" 3

"R" 1 "A" 4

"R" 1 "A" 5

"R" 1 "A" 6

"R" 1 "A" 7

"R" 1 "A" 8

"R" 1 "A" 9

"R" 1 "A" 10

"R" 1 "A" 11

"R" 1 "A" 12

"R" 1 "A" 13

"R" 1 "A" 14

"R" 1 "A" 15

"R" 1 "A" 16

"R" 1 "A" 17

"R" 1 "A" 18

"R" 1 "A" 19

"R" 1 "A" 20

"R" 1 "A" 21

"R" 1 "A" 22

"R" 1 "A" 23

"R" 1 "A" 24

"R" 1 "A" 25

"R" 1 "A" 26

"R" 1 "A" 27

"R" 1 "A" 28

"R" 1 "A" 29

"R" 1 "A" 30

"R" 1 "A" 31

!entry.MVA.unit.name single str

"8CIY.B"

!entry.MVA.unit.positions table dbl x dbl y dbl z

-14.830000 -12.665000 2.038000

-14.885000 -12.102000 3.139000

-13.741000 -12.328000 1.043000

-12.771000 -12.534000 1.471000

-13.861000 -12.920000 0.148000

-13.793000 -11.282000 0.779000

-15.715000 -13.554000 1.734000

-16.817000 -13.940000 2.661000

-16.975000 -15.456000 2.577000

-16.979000 -16.019000 1.492000

-18.124000 -13.171000 2.437000

-17.822000 -11.695000 2.276000

-19.051000 -13.418000 3.618000

-15.714000 -14.267000 0.442000

-16.549000 -13.653000 3.678000

-18.617000 -13.516000 1.528000

-16.863000 -11.574000 1.773000

-18.606000 -11.226000 1.681000

-17.779000 -11.223000 3.258000

-20.015000 -13.773000 3.254000

-18.612000 -14.169000 4.274000

-19.192000 -12.489000 4.171000

-15.194000 -13.667000 -0.304000

-15.207000 -15.225000 0.555000

-16.742000 -14.436000 0.119000

-17.105000 -16.147000 3.658000

-17.255000 -17.590000 3.584000

-17.102000 -15.685000 4.557000

-18.286000 -17.844000 3.388000

-16.640000 -17.986000 2.789000

-16.955000 -18.042000 4.518000

!entry.MVA.unit.residueconnect table  int c1x  int c2x  int c3x  int c4x  int c5x  int c6x

0 0 0 0 0 0

!entry.MVA.unit.residues table  str name  int seq  int childseq  int startatomx  str restype  int imagingx

"ACE" 1 32 1 "?" 0

!entry.MVA.unit.residuesPdbSequenceNumber array int

0

!entry.MVA.unit.solventcap array dbl

-1.000000

0.0

0.0

0.0

0.0

!entry.MVA.unit.velocities table  dbl x  dbl y  dbl z

0.0 0.0 0.0

0.0 0.0 0.0

0.0 0.0 0.0

| | | |
|---|---|---|
| 0.0 | 0.0 | 0.0 |
| 0.0 | 0.0 | 0.0 |
| 0.0 | 0.0 | 0.0 |
| 0.0 | 0.0 | 0.0 |
| 0.0 | 0.0 | 0.0 |
| 0.0 | 0.0 | 0.0 |
| 0.0 | 0.0 | 0.0 |
| 0.0 | 0.0 | 0.0 |
| 0.0 | 0.0 | 0.0 |
| 0.0 | 0.0 | 0.0 |
| 0.0 | 0.0 | 0.0 |
| 0.0 | 0.0 | 0.0 |
| 0.0 | 0.0 | 0.0 |
| 0.0 | 0.0 | 0.0 |
| 0.0 | 0.0 | 0.0 |
| 0.0 | 0.0 | 0.0 |
| 0.0 | 0.0 | 0.0 |
| 0.0 | 0.0 | 0.0 |
| 0.0 | 0.0 | 0.0 |
| 0.0 | 0.0 | 0.0 |
| 0.0 | 0.0 | 0.0 |
| 0.0 | 0.0 | 0.0 |
| 0.0 | 0.0 | 0.0 |
| 0.0 | 0.0 | 0.0 |
| 0.0 | 0.0 | 0.0 |
| 0.0 | 0.0 | 0.0 |
| 0.0 | 0.0 | 0.0 |
| 0.0 | 0.0 | 0.0 |

**Supplementary Note**

**Software Availability and Implementation**

TopFor is implemented as an open-source Python package and is freely available through GitHub (https://github.com/kalininalab/TopFor). The software is distributed as a Git-based repository and can be installed by cloning the source code to a local system.

The core workflow requires Python 3.10 or later, AmberTools (Antechamber, prepgen, parmchk2, and LEaP), and the open-source version of PyMOL; the provided Conda environment uses Python 3.10. The optional RESP charge-derivation workflow additionally requires xTB, ORCA, Multiwfn, and the Python package pexpect. The provided Conda environment installs the core dependencies together with xTB and pexpect, whereas ORCA and Multiwfn must be installed separately for RESP calculations. A setup validation script (setup_env.sh) is provided to configure AMBERHOME and verify the required software components and AMBER parameter files. Detailed installation instructions are provided in the GitHub repository.

TopFor is executed through a unified command-line interface supporting three input modes: single-residue parameterization (topfor -i residue.mol2), whole-peptide parameterization with automatic detection of noncanonical amino acids (topfor -p peptide.mol2), and batch parameterization of multiple residue structures (topfor -b list.txt). Batch input may be provided as a file containing structure paths or as a directory containing the input structures. For single-residue and batch processing, N- or C-terminal residue context can be specified using the --terminal n or --terminal c option, respectively; otherwise, residues are treated as internal. The residue net charge can be specified explicitly using the --net-charge option. The charge-derivation method is selected using the -c option, with AM1-BCC (bcc) as the default and ABCG2 (abcg2), RESP (resp), and Gasteiger (gas) available as alternative charge models.

Comprehensive command-line documentation, example workflows, and source code are available in the GitHub repository (https://github.com/kalininalab/TopFor).

**Output Files and Integration into the AMBER Workflow**

For each processed ncAA, TopFor generates a complete AMBER-compatible residue parameterization package. The output includes charged structural representations (.mol2), residue template files (.prepin and .lib), and force-field modification files (.frcmod), which together define atom types, connectivity, partial charges, and any missing bonded or nonbonded parameters required for molecular simulations. In addition, TopFor generates auxiliary files, including a main-chain definition file (.mc) and JSON metadata describing the residue topology class, head and tail atoms, main-chain atom definitions, terminal context, applied capping groups, selected charge model, final net charge, and any warnings generated during parameterization. These metadata provide a record of the residue-processing and parameterization settings used for each residue.

As part of residue parameterization, TopFor performs a LEaP-based compatibility test. The final MOL2 structure is loaded in LEaP together with the generated protein- and general-force-field modification files (.frcmod), and a library file (.lib) and test topology and coordinate files (.prmtop and .inpcrd) are generated. These test files are used to verify that the parameterized residue and generated force-field parameters can be processed within the AMBER framework and are not the topology and coordinate files used for subsequent peptide or protein–peptide simulations. For downstream applications, the TopFor-generated residue template (.prepin) and selected force-field modification file or files can be loaded separately in LEaP during preparation of the complete peptide or protein–peptide system. A representative output directory generated for a single residue is shown below:

```
<output_base>/
├── MVA.mol2
├── MVA.split.json
└── MVA/
    ├── residue.mol2
    ├── residue_capped.mol2
    ├── residue_capping_meta.json
    ├── residue_meta.json
    ├── MVA.mol2
    ├── MVA.ac
    ├── MVA.mc
    ├── MVA.prepin
    ├── MVA.lib
    ├── MVA_ff19SB.frcmod
    ├── MVA_gaff2.frcmod
    ├── leap.in
    ├── MVA.log
    ├── MVA.inpcrd
    └── MVA.prmtop
```

Completed and failed TopFor runs are additionally recorded in summary text files to facilitate batch processing and identification of residues requiring further inspection.

#### Supplementary Figures

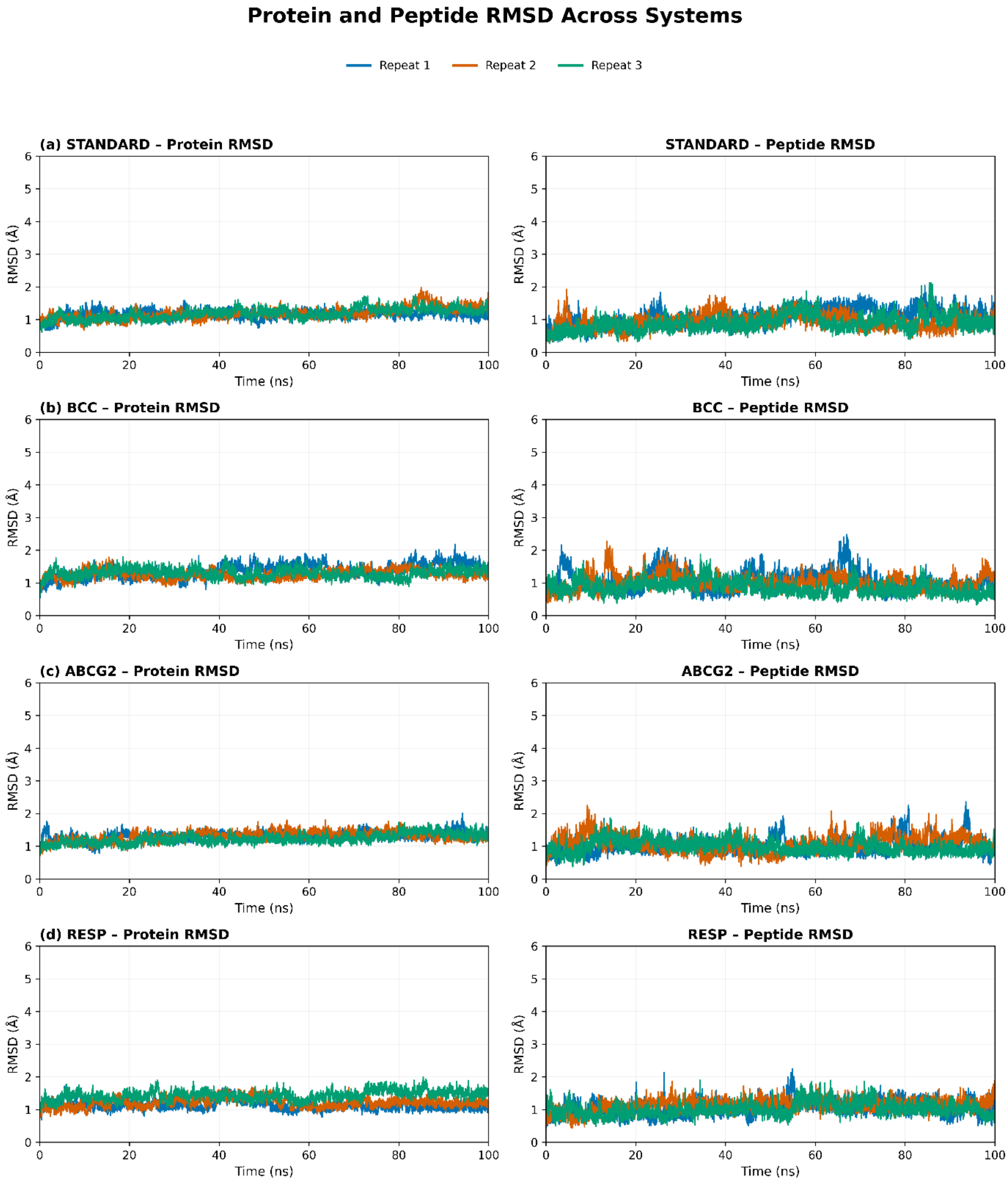

**Figure S1. Backbone RMSD for the MDM2–p53 validation system** during three independent 100 ns production simulations. RMSD profiles are shown for the STANDARD (a), AM1-BCC (b), ABCG2 (c), and RESP (d) parameterization schemes, with the MDM2 receptor and bound p53 peptide shown separately. The three independent simulations are represented by different colors in each panel. Similar RMSD profiles across the parameterization schemes indicate comparable structural stability of the MDM2–p53 complexes.

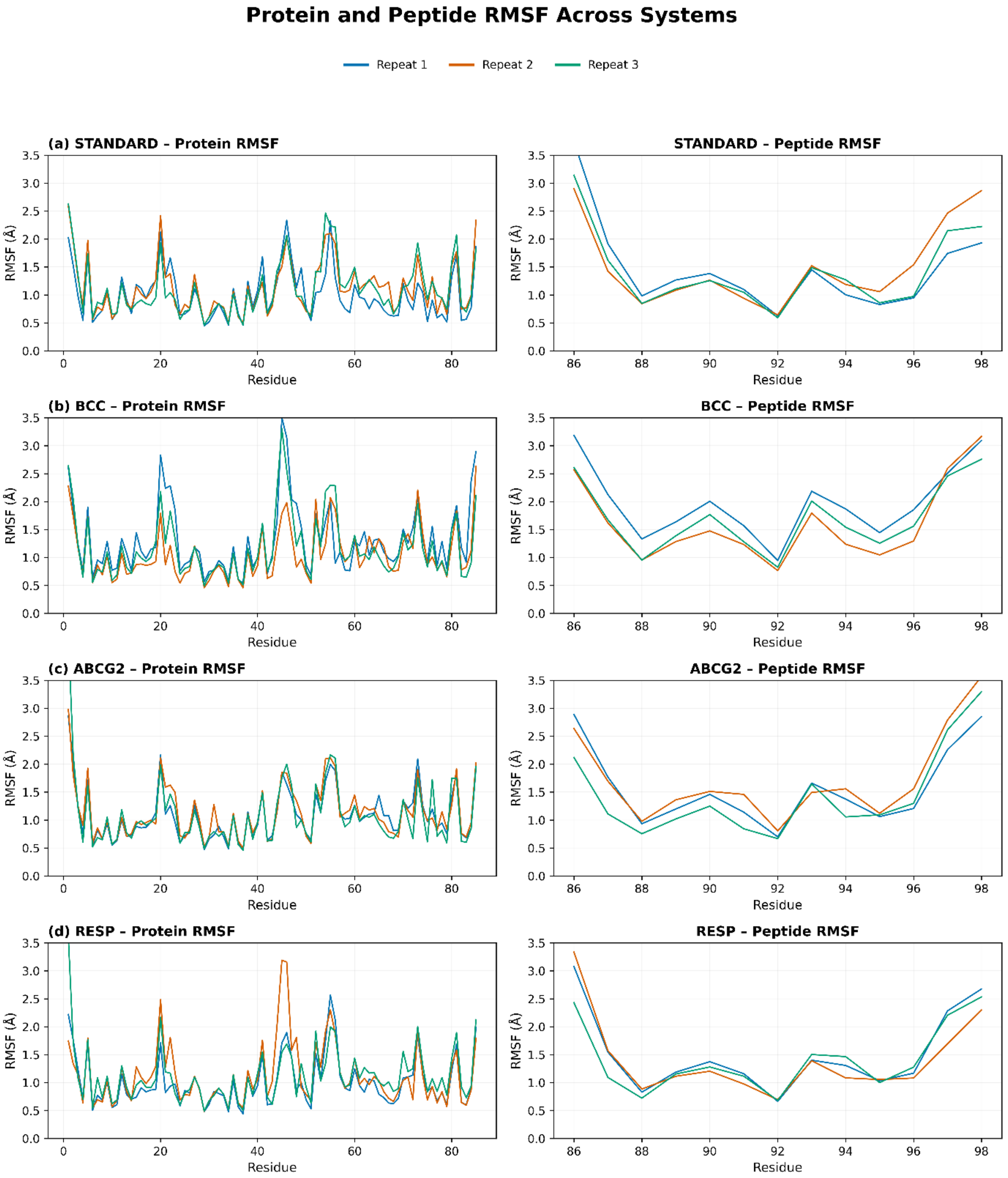

**Figure S2. Per-residue Cα RMSF for the MDM2–p53 validation system** during three independent 100 ns production simulations. RMSF profiles are shown for the STANDARD (a), AM1-BCC (b), ABCG2 (c), and RESP (d) parameterization schemes and report residue-level flexibility of the MDM2 receptor and bound p53 peptide. The three independent simulations are represented by different colors in each panel. Similar RMSF profiles across the parameterization schemes indicate comparable residue-level flexibility.

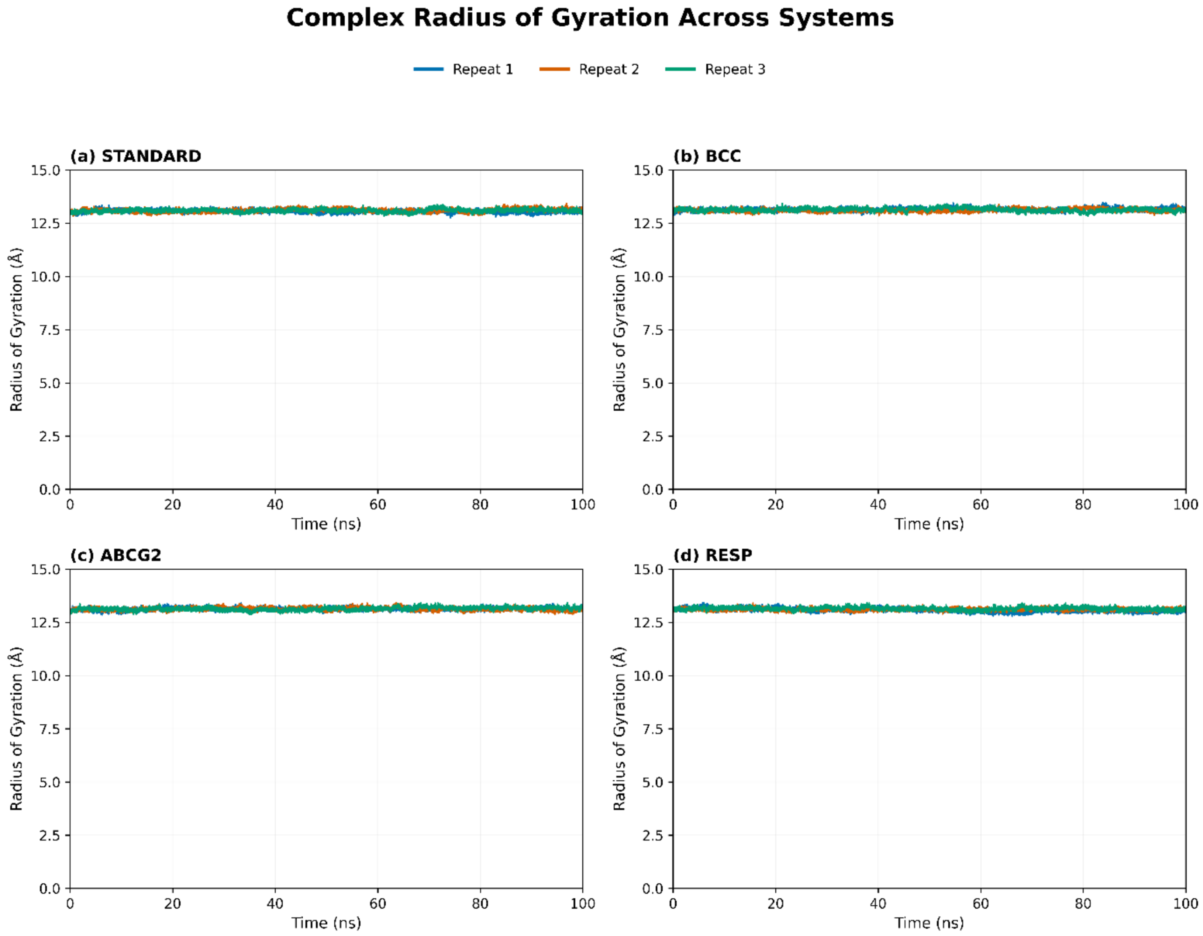

**Figure S3. Radius of gyration (Rg) of the MDM2–p53 complex** during three independent 100 ns production simulations. Rg profiles are shown for the STANDARD (a), AM1-BCC (b), ABCG2 (c), and RESP (d) parameterization schemes. The three independent simulations are represented by different colors in each panel. Comparable Rg profiles across the parameterization schemes indicate preservation of the global compactness of the MDM2–p53 complex.

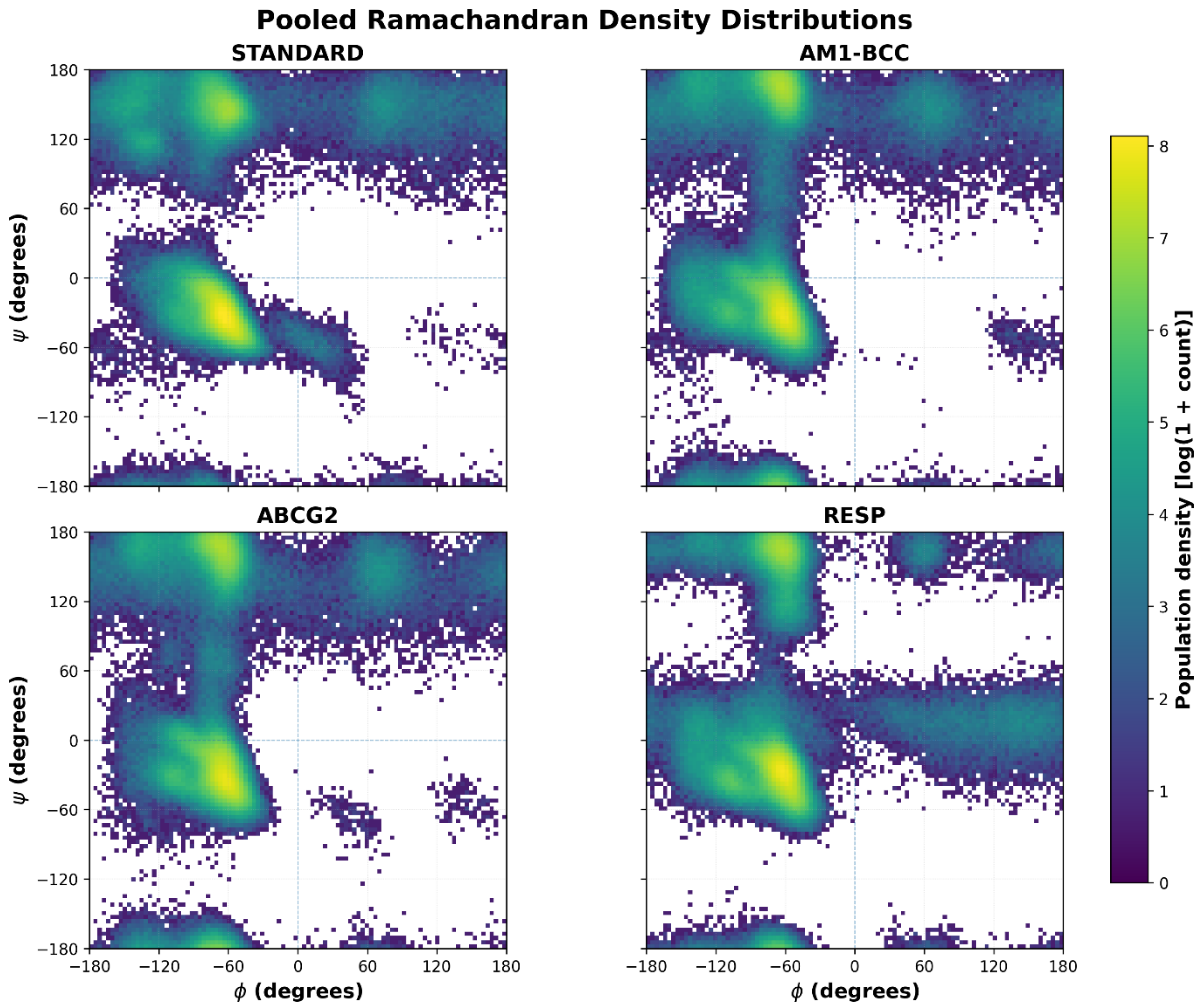

**Figure S4. Pooled backbone Ramachandran density distributions of the p53 peptide** for the four parameterization schemes. The distributions were generated by combining all frames from the three independent 100 ns production trajectories for each parameterization scheme (STANDARD, AM1-BCC, ABCG2, and RESP). Color intensity represents the frequency of sampling, with darker colors indicating sparsely populated backbone conformations and brighter green-to-yellow regions corresponding to increasingly populated ϕ/ψ conformational states. The highly similar distributions indicate comparable backbone conformational sampling across the four parameterization schemes.

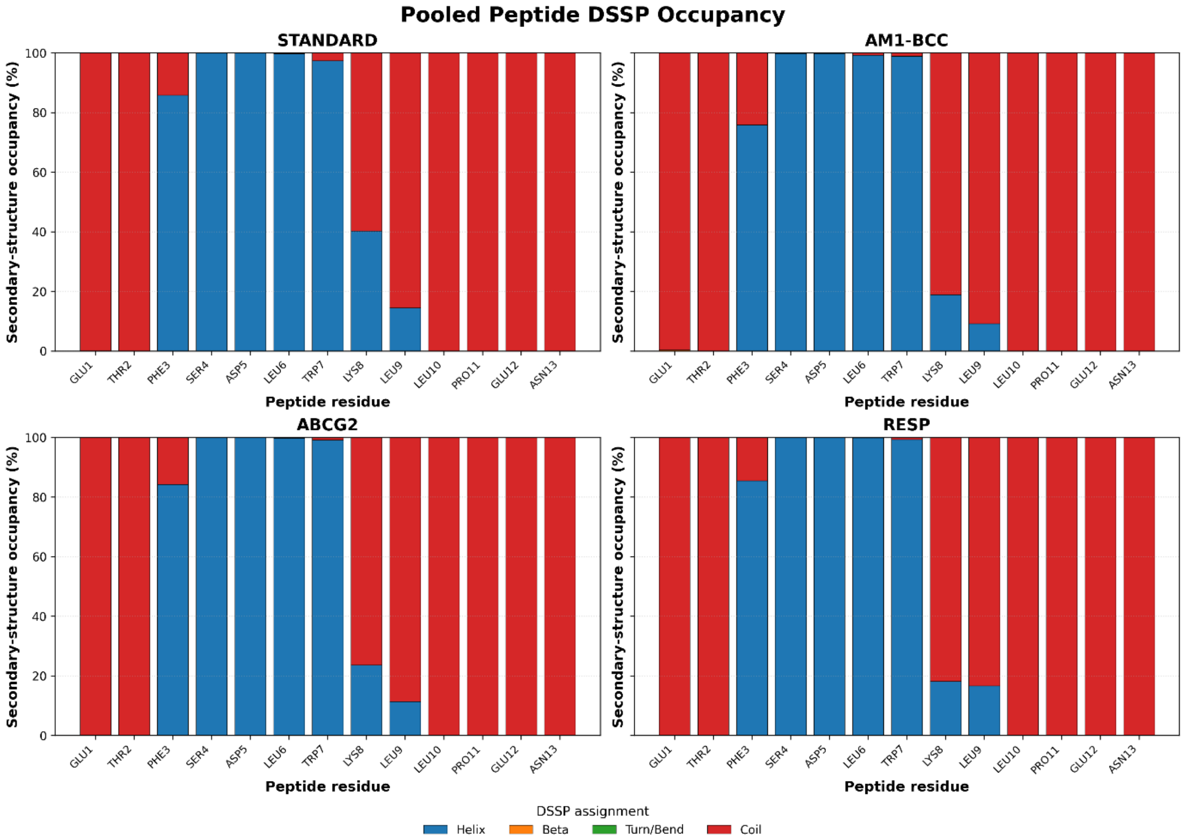

**Figure S5. Pooled DSSP secondary-structure occupancies of the p53 peptide** for the four parameterization schemes. Secondary-structure assignments were combined from the three independent 100 ns production trajectories for each parameterization scheme (STANDARD, AM1-BCC, ABCG2, and RESP). The central residues consistently maintained the crystallographic α-helical conformation, whereas the terminal residues predominantly adopted coil conformations, indicating preservation of the native secondary-structure profile across all TopFor-generated parameterizations.

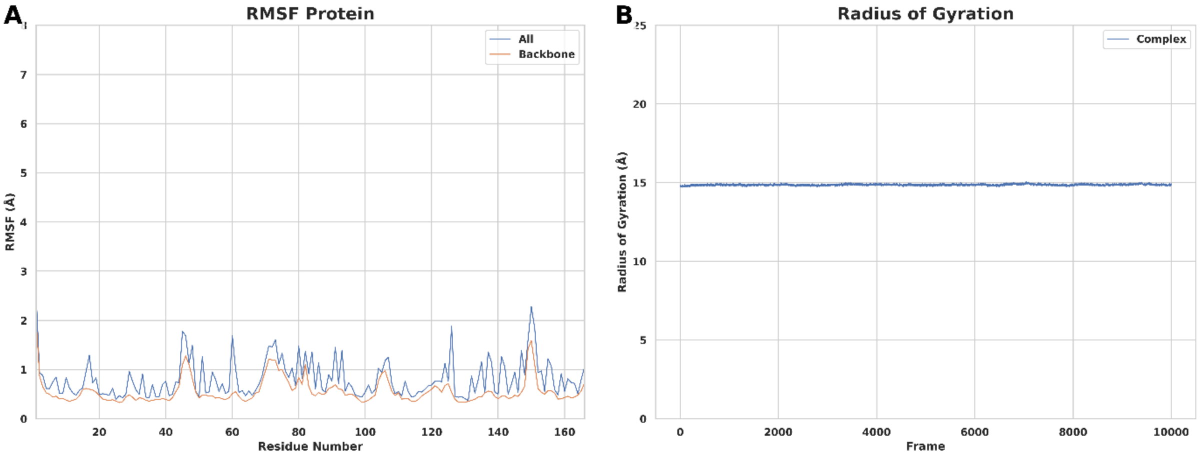

**Figure S6. Additional molecular dynamics analyses of the cyclophilin–cyclosporin A complex (PDB ID: 1XQ7).** (A) Cα RMSF of cyclophilin. (B) Radius of gyration (Rg) of the cyclophilin–cyclosporin A complex during the 100 ns production simulation.

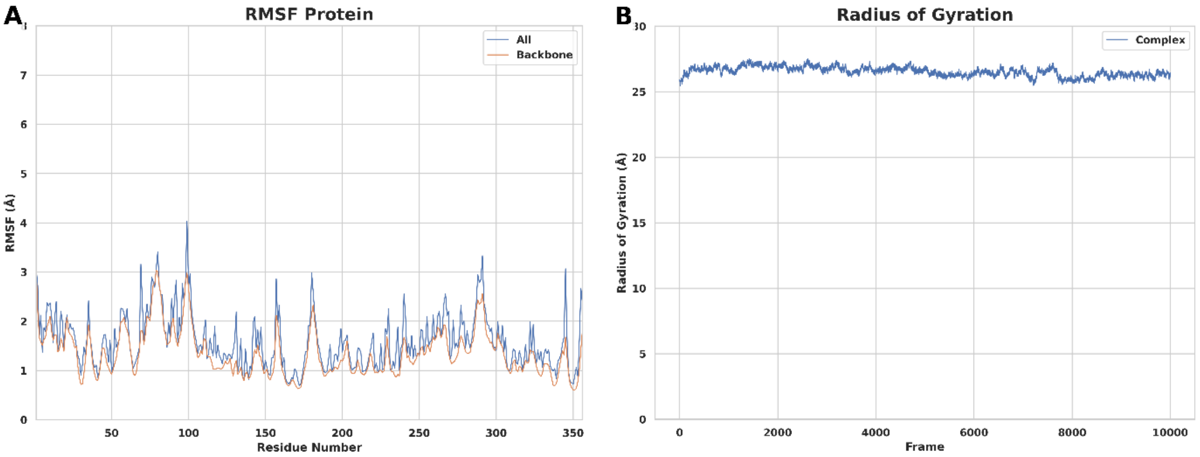

**Figure S7. Additional molecular dynamics analyses of the native DnaN–cyclohexylgriselimycin (CGM) complex (PDB ID: 8CIY). (A) Cα RMSF of CGM. (B) Radius of gyration (Rg) of the** DnaN–cyclohexylgriselimycin complex during the 100 ns production simulation.

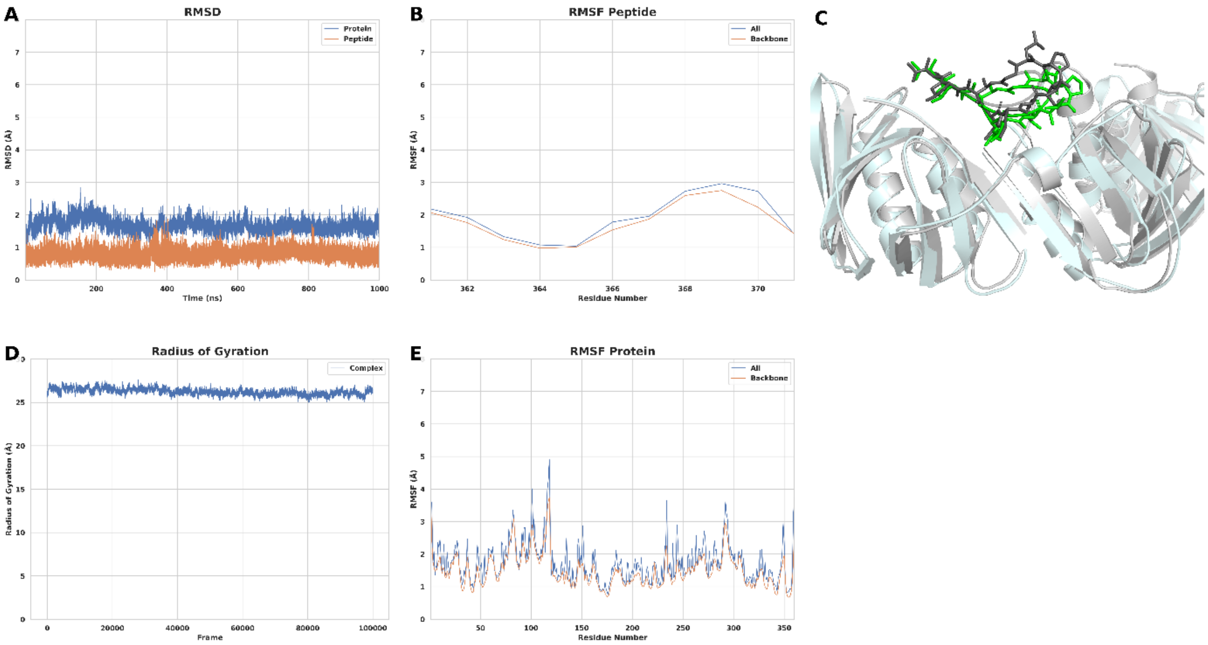

**Figure S8. Molecular dynamics simulation results of the *Escherichia coli* DnaN–griselimycin (GM) complex (PDB ID: 8CIX).** (A) Backbone RMSD of DnaN and the bound GM peptide during the 1000 ns production simulation. (B) Cα RMSF of the bound GM peptide. (C) Superposition of the representative structure obtained by clustering the 1000 ns production trajectory with the experimental crystal structure. DnaN is shown as a cartoon, with the crystal structure in light gray and the MD representative structure in pale cyan. GM is shown as sticks, with the crystal structure in gray and the MD representative structure in green. (D) Radius of gyration (Rg) of the DnaN–GM complex during the 1000 ns production simulation. (E) Cα RMSF of DnaN.

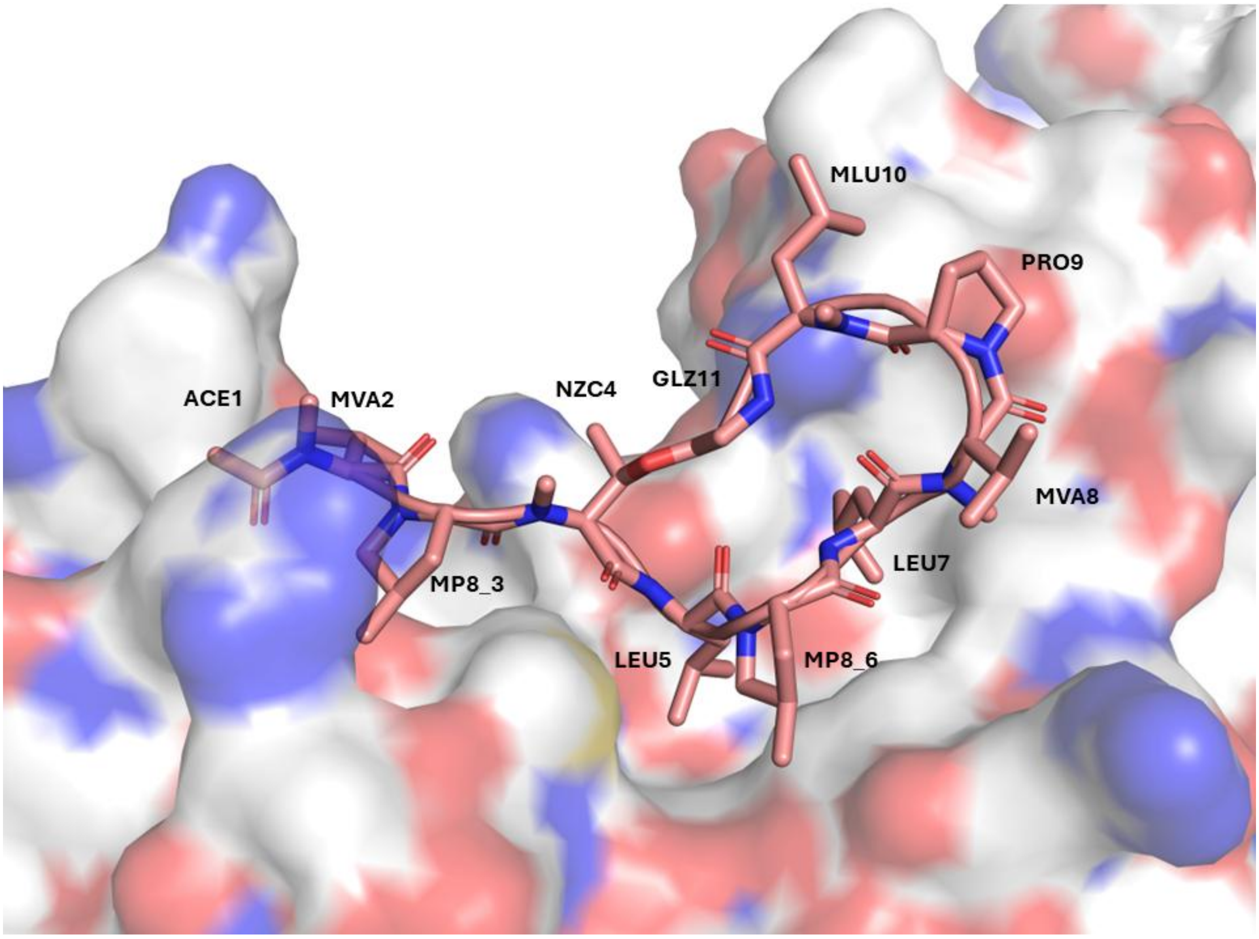

**Figure S9.** ***Escherichia coli*** **DnaN–GM complex (PDB ID: 8CIX).** The protein is shown as a surface representation, with negatively charged residues colored red, positively charged residues blue, and neutral residues white. GM is shown as sticks, with carbon atoms colored salmon, nitrogen atoms blue, and oxygen atoms red. The amino acid residues of GM are labeled.

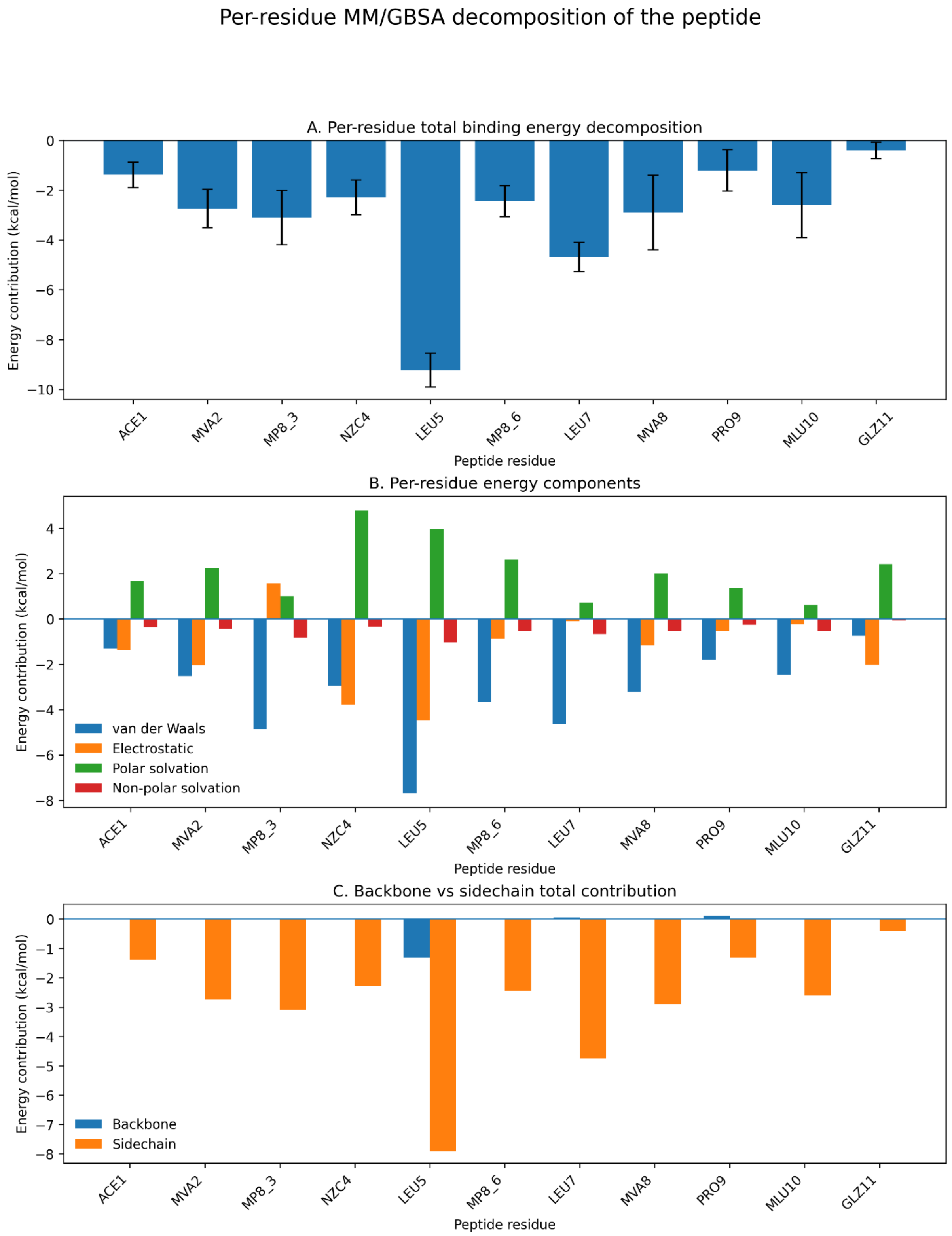

**Figure S10. MM/GBSA per-residue energy decomposition analysis of GM in the DnaN–GM complex (PDB ID: 8CIX).** (A) Per-residue total binding energy contributions. (B) Per-residue contributions of individual energy components, including van der Waals, electrostatic, polar solvation, and nonpolar solvation energies. (C) Backbone and side-chain contributions to the total binding energy.

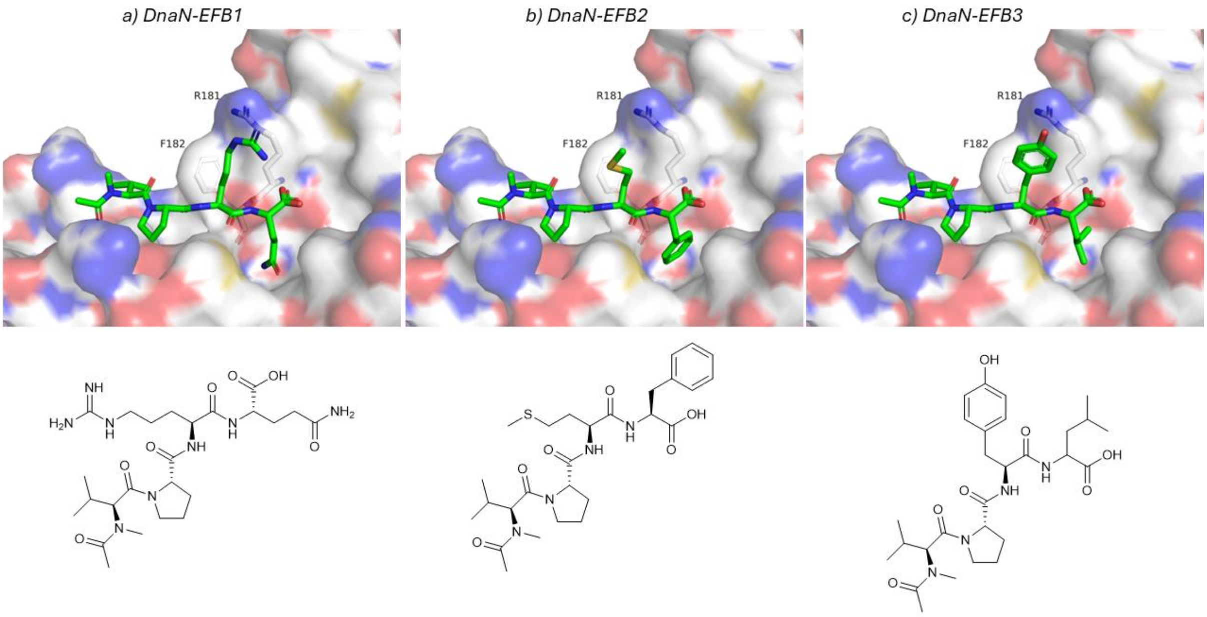

**Figure S11. Docking poses of EFB1–EFB3 in *Mycobacterium smegmatis* DnaN (PDB ID: 5AH2).** The protein is shown as a surface representation, with negatively charged residues colored red, positively charged residues blue, and neutral residues white. EFB1–EFB3 are shown as sticks, with carbon atoms colored green, nitrogen atoms blue, and oxygen atoms red. The corresponding 2D structures of EFB1–EFB3 are shown below the docking poses.
